# The *Drosophila* Dosage Compensation Complex activates target genes by chromosome looping within the active compartment

**DOI:** 10.1101/101634

**Authors:** Tamás Schauer, Yad Ghavi-Helm, Tom Sexton, Christian Albig, Catherine Regnard, Giacomo Cavalli, Eileen E M Furlong, Peter B Becker

**Affiliations:** Molecular Biology Division, Biomedical Center and Center for integrated Protein Science Ludwig-Maximilians-University, Munich, Germany; European Molecular Biology Laboratory, Genome Biology Unit, Heidelberg, Germany; Institute of Genetics and Molecular and Cellular Biology, Illkirch, France; Institute of Human Genetics, CNRS, Montpellier, France; University of Montpellier, France

**Keywords:** *Drosophila* dosage compensation, nuclear architecture, chromosome conformation capture

## Abstract

X chromosome dosage compensation in *Drosophila* requires chromosome-wide coordination of gene activation. The male-specific-lethal dosage compensation complex (DCC) identifies X chromosomal High Affinity Sites (HAS) from which it boosts transcription. A sub-class of HAS, PionX sites, represent first contacts on the X. Here, we explored the chromosomal interactions of representative PionX sites by high-resolution 4C and determined the global chromosome conformation by Hi-C in sex-sorted embryos. Male and female X chromosomes display similar nuclear architecture, concordant with clustered, constitutively active genes. PionX sites, like HAS, are evenly distributed in the active compartment and engage in short- and long-range interactions beyond compartment boundaries. By *de novo* induction of DCC in female cells, we monitored the extent of activation surrounding PionX sites. This revealed a remarkable range of DCC action not only in linear proximity, but also at megabase distance if close in space, suggesting that DCC profits from pre-existing chromosome folding to activate genes.

## Introduction

The process of dosage compensation in *Drosophila melanogaster* involves doubling productive transcription of essentially all active genes on the single male X chromosome to match the combined output of the two female Xs. It provides an instructive example of coordinated, chromosome-wide regulation of transcription (for a review see (Lucchesi & Kuroda 2015; Keller & Akhtar 2015)). The regulator of this process, the male-specific-lethal dosage compensation complex (MSL-DCC or just DCC) consists of five protein subunits (MSL1, MSL2, MSL3, MOF and MLE) and two long, non-coding RNAs (*roX1* and *roX2*). Targeting the DCC to the X chromosome is thought to occur in at least two steps. The complex first recognizes and binds to about 250 so-called High Affinity Sites (HAS; also referred to as ‘chromosomal entry sites, CES), from which it ‘spreads’ by an unknown mechanism to acetylate the chromatin of active genes in its vicinity (Alekseyenko et al. 2008; Straub et al. 2008; Alekseyenko et al. 2012) (Straub et al. 2013; Ferrari et al. 2013). HAS often contain a GA-rich, low-complexity sequence motif, the MSL Recognition Element (MRE). We recently found that MSL2, the only male-specific protein subunit of the DCC, is able to bind MREs with high selectivity (Villa et al. 2016). This study also revealed that MSL2 uses its CXC domain to identify a distinct sequence motif characterized by a crucial extension of the classical MRE signature, which are highly enriched on the X chromosome. These sites, which we termed ‘PionX’ sites (pioneering sites on the X), are characterized by a novel DNA conformation signature (Villa et al. 2016). Sites bearing the PionX signature are the first contacts of MSL2 upon *de novo* induction of DCC assembly in female cells. The mechanism through which the initial binding of DCC to a fairly small number of PionX sites (56 have been defined) initiates the activation of all genes on the X remain unknown. Conceivably, propagation of the DCC’s H4K16 acetylation activity may profit from the folding of the X chromosome.

‘Chromosome conformation capture’ technologies, as well as DNA FISH, revealed that chromosomes are organized at different scales. At the highest level chromosomal domains are partitioned into active (A) and inactive (B) compartments, which alternate when mapped onto the linear chromosome, but may cluster in space to organize the interphase nucleus (Lieberman-Aiden et al. 2009; Dekker et al. 2013; Sexton & Cavalli 2015). Changes in cell type-specific gene activity during differentiation and the associated epigenetic marks correlate with corresponding changes of compartment patterns (Dixon et al. 2015; Fortin & Hansen 2015). Within each compartment, regions of increased chromosomal interactions can be visualized, termed ‘topologically associated domains’ (TADs). TADs may be thought of as functional units of the hierarchical genome organization, since their positions and boundaries are invariant across cell-types in mammals (Nora et al. 2012; Dixon et al. 2012) and in *Drosophila* (Ramírez et al. 2015) (Ulianov et al. 2016). The appearance of TADs in chromosome conformation mapping may reflect insulator interactions or interactions between chromatin segments mediated by multivalent structural proteins (Dixon et al. 2012; Sexton et al. 2012; Rowley & Corces 2016). Sub-TAD interactions generally reflect looping interaction of *cis*-regulatory elements at active loci (Ghavi-Helm et al. 2014).

The function of the DCC might contribute to or profit from the three-dimensional chromatin organization at any of these levels. The DCC interacts with HAS and nucleosomes in transcribed genes through distinct surfaces and we hypothesized that the complex may ‘scan’ the proximal space for target chromatin while bound to HAS (Straub et al. 2013). The TAD organization of the *Drosophila* X chromosome has been characterized in detail and was found invariant in cell lines with male or female karyotype and not dependent on active dosage compensation (Ramírez et al. 2015; Ulianov et al. 2016).

Given the prominent role of PionX sites in DCC recruitment we explored the relationship between representative PionX sites and chromosome architecture using both high-resolution 4C and Hi-C, which we derived from sex-sorted *Drosophila* embryos. An analysis at the level of the chromosomal compartments, which are poorly studied in *Drosophila* embryos, was very informative. We found that acetylation of histone H4 at lysine 16 (H4K16ac) serves well to mark the active compartment. Despite quantitative differences in H4K16ac levels at male and female X chromosomes, the overall chromosome structure is invariant. PionX and other HAS distribute evenly throughout the active compartment in proximity to clustered active genes. High-resolution 4C-seq experiments showed that PionX loci are in spatial contact with many other loci within the active compartment, with no specific requirement for HAS at target sites. Remarkably, these interactions frequently span across large regions of inactive compartments, suggesting that they are enabled by spatial clustering of active domains. To more quantitatively assess the extent of DCC action, we induced *de novo* DCC binding to a small number of PionX sites in female cells, revealing that the DCC activates target genes by chromosome looping.

## Results

### The X chromosome compartment architecture is similar in male and female embryos

Previous Hi-C data for *Drosophila* were generated either in mixed-sex embryos (Sexton et al. 2012) or in cultured cells with known sex (Ramírez et al. 2015) (Ulianov et al. 2016). These cell lines may not be directly comparable to each other for sex- and dosage compensation-related differences, because they derive from different tissues, bear large genomic rearrangements (Ramírez et al. 2015) or differ greatly in genomic copy number (Lee et al. 2014). To better assess potential sex-specific chromosome conformation differences we sorted 16-18 h embryos according to sex and generated Hi-C libraries (**Supplementary File 1**).

We created coverage-normalized contact enrichment maps as well as coverage- and distance-normalized correlation maps with 10 kb resolution (Figure 1A,B and Figure 1 – figure supplement 1A; and see Methods (Heinz et al. 2010)). Both normalization approaches reveal very similar contact patterns for the two sexes. To determine the compartment architecture of the genome, we performed principal component analysis (PCA) on the correlation matrix (Lieberman-Aiden et al. 2009). The first principal component (PC1, also called eigenvector) demarcates domains in the A and B compartment (Figure 1C; Figure 1 – figure supplements 1B,C and **Supplementary File 2**). We define the sign of the PC1 vector such that regions with positive values are enriched for peaks of acetylation of histone H4 at lysine 16 (H4K16ac), and hence are referred to as active chromatin (**also see below**). Accordingly, we call the compartment with negative PC1 value ‘inactive’.

**Figure 1.**
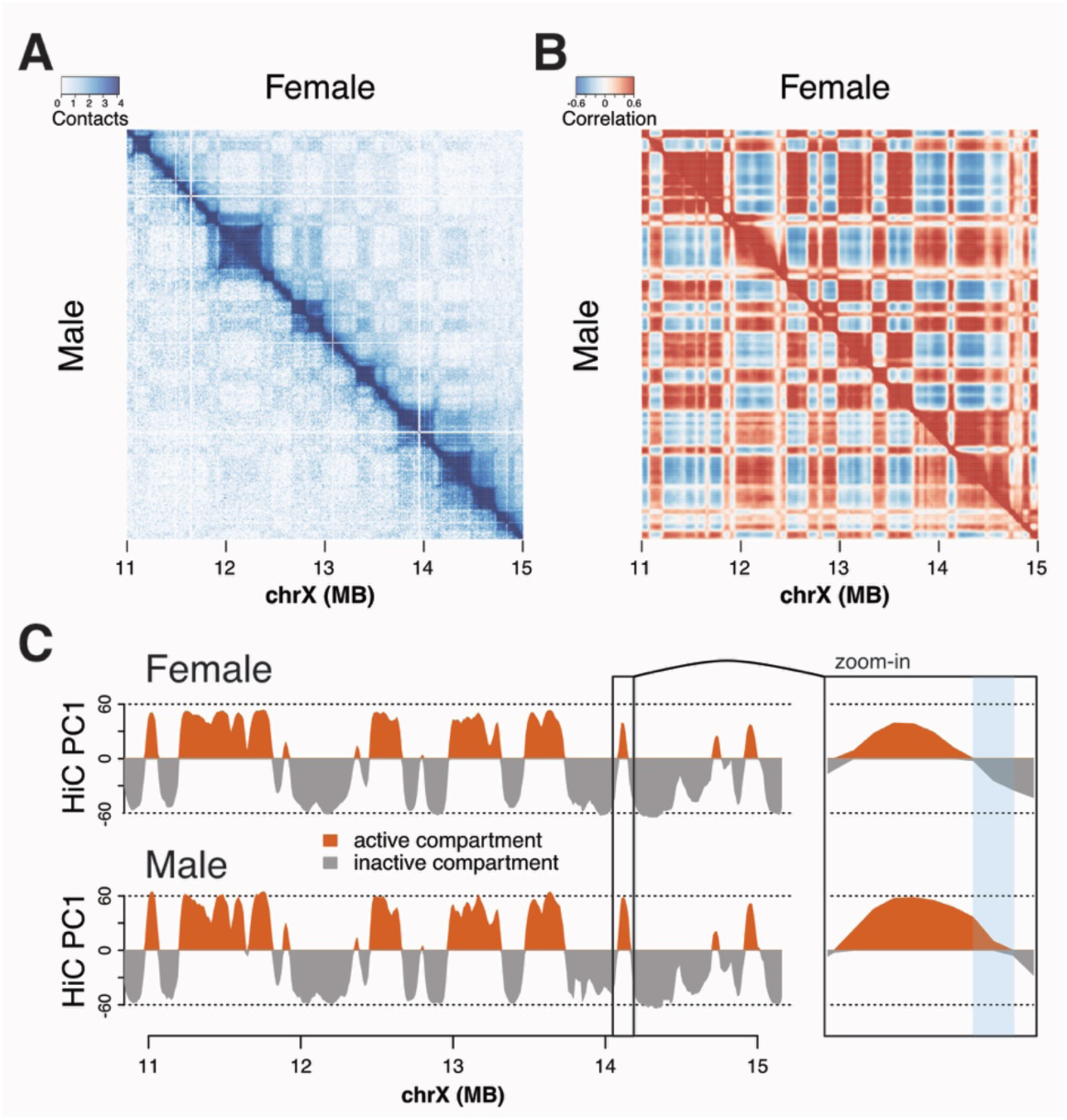
Hi-C contacts maps derived from male and female embryos reveal similar threedimensional topology of the X chromosome. **A**) Coverage-normalized Hi-C contact maps of the X chromosome at the region 11-15 MB in female (*upper triangular matrix*) and male (*lower triangular matrix*) embryos (resolution: 10kb). **B**) Smoothed, coverage- and distance-normalized correlation matrix of the same region as in A) (color key: red - positive, blue - negative correlation, resolution: 10kb with 50 kb smoothing). **C**) First principal component (PC1) of the correlation matrix in B). Positive values (red) indicate the active and negative values (grey) the inactive compartment. Right frame: Zoom-in view of a locus switching compartments between female and male. See PC1 values in Supplementary File 2.

To further characterize the compartment architecture we defined continuous regions (domains) within the active or inactive compartment. The size distribution of active and inactive domains is in the same range with medians around 100 kb on the X and 2L chromosomes in both sexes. Larger domains tend to be more loosely organized, judged by the inverse relationship between domain size and average contact enrichment (Figure 1 – figure supplement 2A). Independent of size, the contact enrichment values for inactive domains are always higher than those of active domains, as expected (Ulianov et al. 2016) (Figure 1 – figure supplement 2A,B). This is illustrated by the steeper intra-domain off-diagonal signal decay for active compared to inactive domains (Figure 1 – figure supplement 2C).

In summary, inactive domains are more compact and homogenous compared to looser, active domains and these characteristics do not differ either between females and males or between chromosome X and 2L.

### Compartment switching correlates with differential marking of H4K16ac between females and males

Correlation of PC1 between sexes and biological replicates was remarkably high on the X chromosome as well as on chromosome 2L (Pearson´s r = 0.96-0.99; Figure 1 – figure supplements 1B). However, the PC1 varied more between females and males on the X chromosome compared to chromosome 2L and between replicates (Figure 1 – figure supplement 1B. We also note that the positive PC1 values are 15-20% higher on the male X compared to female suggesting enhanced interactions throughout the active compartment on this chromosome (Figure 1C and Figure 1 – figure supplement 1B). Beside the overall difference in PC1 values, 7-8% of the X-chromosomal fragments switch sign between sexes, but only 2-3% between replicates, suggesting that the corresponding loci reside in different compartments in female and male embryos (p-value < 10^-7^, Fisher´s exact test; Figure 1C and Figure 1 – figure supplement 1C).

To verify these differences using another approach, we calculated the correlation on the contact profiles between the two sexes for each locus one-by-one and displayed them along the genome. The median correlation was relatively high for both chromosomes (chrX: r ~0.93; chr2L: r ~0.98) however, 15% of the fragments on the X chromosome have a lower than 0.8 correlation of contact enrichment, while this is not the case for chr2L (Figure 1 – figure supplement 1C, upper panel). A lower median correlation (r ~0.75) characterizes loci that reside in different compartments in male and female nuclei.

To investigate whether these differences can be related to chromatin marks as suggested in human cells (Fortin & Hansen 2015); (Dixon et al. 2015) we compared the female or male PC1 values to two histone modifications that mark actively transcribed chromatin, H3K36me3 and H4K16ac, and to gene expression (i.e. RNA-seq) (Filion et al. 2010); (Prestel et al. 2010) in Kc (female) and S2 (male) cells (Figure 1 – figure supplement 3). Fragments with positive PC1 have higher levels of H4K16ac, H3K36me3 and RNA-seq signal in both cells, which confirms their residency in the active compartment. Hi-C fragments have higher average H4K16ac levels on the X chromosome in S2 cells as expected from the known role of this modification at gene bodies during dosage compensation (Prestel et al. 2010); (Lucchesi & Kuroda 2015); (Kind et al. 2008) (Figure 1 – figure supplement 3A). Interestingly, Hi-C fragments that were marked by increased H4K16ac also have higher PC1 values in males suggesting a correlation between three-dimensional chromatin interactions and the action of the DCC on target gene chromatin (Figure 1 – figure supplement 3A). Zooming in to these loci reveals that in several cases the compartment boundaries are shifted such that the active H4K16ac domain is extended at the expense of the inactive compartment (Figure 1 – figure supplement 4). Curiously, the reverse was not the case: fragments of active compartment in females that reside in the inactive compartment in males did not systematically differ in their H4K16ac marks (Figure 1 – figure supplement 3A). In both scenarios the H3K36me3 and RNA-seq levels were unchanged illustrating that the enhanced in PC1 on the single male X accompanies dosage compensation (Figure 1 – figure supplement 3B,C and 4).

In summary, we found that the overall architecture of the X chromosome is similar in male and female embryos, and similar to cell lines of male and female genotype, in line with previous observations (Ramírez et al. 2015). In addition, we observe that X chromosomal fragments differ slightly in their three-dimensional contacts (i.e. PC1) in male and female embryos and correlate with H4K16ac levels.

### Hallmarks of the dosage-compensated compartment on the male X

Our compartment definition reveals that epigenetic and topological domains overlap highly in *Drosophila*, in agreement with previous findings (Sexton et al. 2012). For example, domains of constitutive transcription (“yellow” chromatin (Filion et al. 2010)) essentially always localize to the active compartment (Figure 2 – figure supplement 1A,B). New RNA-seq data obtained from S2 and Kc cells confirm that genes located in the active compartment have higher expression compared to the inactive compartment (Figure 2 – figure supplement 1C). The DCC acts on actively transcribed chromatin, which is enriched in H3K36 methylation, a hallmark of ‘yellow’ chromatin (Larschan et al. 2007; Sural et al. 2008; Filion et al. 2010). The RNA-seq data also confirm the commonplace notion that disrupting DCC in S2 cells by MSL2 RNAi or inducing DCC in Kc cells by SXL RNAi preferentially modifies gene expression levels in the active compartment and does not change transcription of inactive genes or autosomal genes (Alekseyenko et al. 2012) data (Figure 2 – figure supplement 1D).

**Figure 2.**
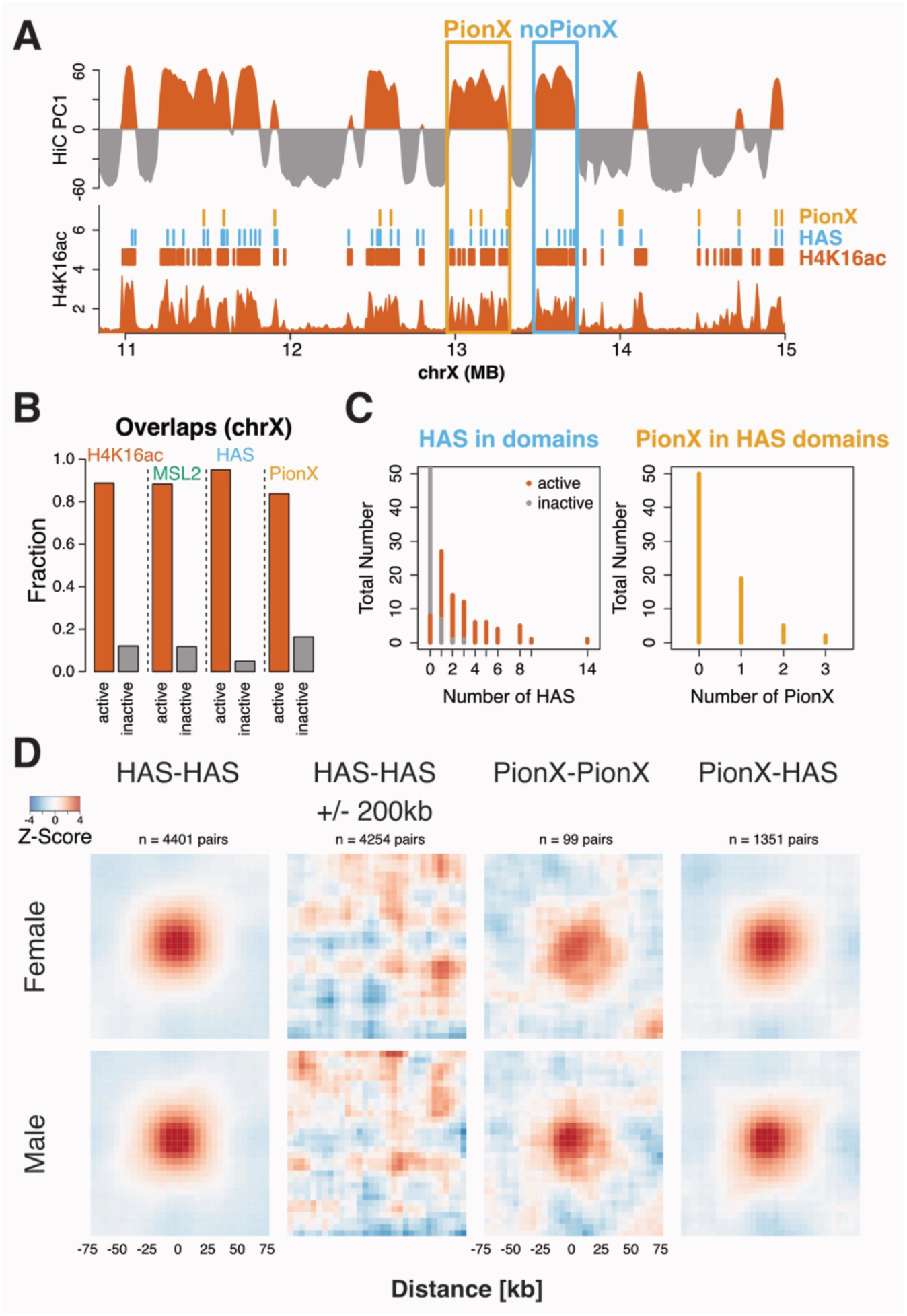
Features of dosage compensation localize to the active compartment. **A**) *Top track*: First principal component (PC1) of the correlation matrix in male embryos (as in Figure 1). *Bottom track*: H4K16ac ChIP-seq signal along the same region in S2 cells (male). PionX sites, High Affinity sites (HAS) and H4K16ac peak regions are indicated with small bars. Examples of domains with or without PionX sites are framed. **B**) Fraction of H4K16ac, MSL2, HAS and PionX sites overlapping domains located in the active (red) and inactive (grey) compartment. **C**) *Left*: Number of HAS within domains located in the active (red) and inactive (grey) compartment. *Right*: Number of PionX sites within domains containing HAS in the active compartment (ocher). **D**) Inter-domain average contact enrichment sub-matrices centered around pairs of HAS, control sites (HAS +/- 200 kb), PionX sites and PionX-HAS in females (*top*) and males (*bottom*). Interactions between 250 kB and 2.5 MB distance are considered, resolution: 5kb with 25 kb smoothing.

The concept that the DCC acts only in restricted parts of the genome is further supported by the finding that the vast majority of MSL2 binding sites, HAS and PionX sites, reside in the active compartment (Figure 2A,B and Figure 2 – figure supplement 2A). There they appear roughly evenly distributed, so that larger domains contain more HAS (Figure 2 – figure supplement 2B). Seventy-six out of 84 active domains contain at least one, on average three HAS (Figure 2C). However, not every active domain contains a PionX site, in fact the majority (50) does not contain any (Figure 2A,C, Figure 2 – figure supplement 2A). This reveals that DCC action at target genes does not require a PionX site within each active domain. Domains with PionX sites tend to be larger than domains with only ‘regular’ HAS and, consequently, have less intra-domain contact enrichments (Figure 2 – figure supplement 2C).

Many loci in the *Drosophila* genome, such as polycomb response elements or enhancers, are engaged in long-range, inter-domain interactions (Sexton et al. 2012; Ghavi-Helm et al. 2014). It has been shown that HAS are particularly engaged in long-range contacts in various cell lines (Ramírez et al. 2015). We find that similar levels of pair-wise interactions between HAS (including PionX sites) in a range of 250 kb – 2.5 Mb in female and male embryos (Figure 2D and Figure 2 – figure supplement 3A). Loci at distances +/-200 kb away from HAS do not show an enrichment, confirming the special engagement of HAS at this resolution (Figure 2D). Interestingly, inter-domain interactions between PionX sites are slightly stronger (on average 1.5x) and more focused in males compared to females (Figure 2D). Iterative resampling of sites (30 sites, 1000 iteration) confirms the increase in long range contact enrichment between PionX sites, while ‘ordinary’ HAS show no sex-specific interaction bias (Figure 2 – figure supplement 3A). As HAS are mainly located in the active chromatin we also investigated these contacts in the ‘yellow’ chromatin domain and find similar pattern as for HAS (Figure 2 – figure supplement 3B) suggesting that long-range interactions are a general feature of active chromatin.

Taken together, the DCC resides almost exclusively in the active nuclear compartment that constitutes an environment permissive for long-range interactions. Long-range interactions between PionX sites are stronger in males, revealing a tendency to cluster locally in the nucleus.

### High-resolution 4C-seq reveals interactions between PionX sites and active domains

In order to define the chromosomal interactions of selected PionX sites, we generated high-resolution profiles by 4C-seq in S2 cells (Ghavi-Helm et al. 2014; Klein et al. 2015)(**Supplementary File 4-6**). We chose viewpoints that are either very prominent PionX sites and/or sites that are among the first to be bound by MSL2 *in vivo*, if dosage compensation is induced in female cells (Villa et al. 2016) (**Supplementary File 7**). Female Kc cells repress MSL2 expression and thus the male phenotype through the regulator sex lethal (SXL). If SXL is depleted by RNA interference, MSL2 and *roX* expression ensues, the DCC forms and activates genes in the active compartment (Figure 2 – figure supplement 1D) (Alekseyenko et al. 2012). Reprogramming cells in this way takes several days during which one can map the very first chromosomal contacts of the complex, which bear PionX signature (Villa et al. 2016). We refer to these sites as ‘induced’ MSL2 binding sites in the later analysis.

4C profiles at selected viewpoints in cell lines were remarkably similar to virtual 4C profiles extracted from the Hi-C contact maps for the same loci in embryos (median correlation: ~0.65; Figure 3A and Figure 3 – figure supplement 1A,B). We monitored the interaction network of these viewpoints and found them engaged in a total of 805 interactions (peaks) with an average of 73 interactions per viewpoint (**Supplementary File 5**). As expected, most of the interactions (~80%) were located in the active compartment (Figure 3B). An exceptional case is the viewpoint at ~19.9 MB that resides in the inactive compartment and makes restricted contacts to active and inactive domains surroundings (Figure 3B and Figure 3 – figure supplement 1). The distance between viewpoints and interaction peaks varies greatly with a median of 500 kb including long-range interactions reaching up to 10 Mb (Figure 3 – figure supplement 2A). Those viewpoints that contact fewer and only closer regions do not reside in a typical, large active domain but are either close to the boundary (3.8 Mb viewpoint) or in a very small active domain surrounded by inactive domains (12.8 MB viewpoint) or located within an inactive domain (19.9 MB viewpoint) (Figure 3 and Figure 3 – figure supplement 1). All other viewpoints reside well within active compartments, and their chromosomal contacts are likewise located in active domains (i.e. ‘yellow’ chromatin) and also marked by H4K16ac (Figure 3 – Figure 3 figure supplement 2B,C). PionX sites do not preferentially contact other HAS. The 4C interaction peaks overlap only moderately with MSL2 bindings sites (55%) and even less so with HAS (~25%) (Figure 3C and Figure 3 – figure supplement 2C) indicating that the viewpoints are in contact with entire domains and not individual sites.

**Figure 3.**
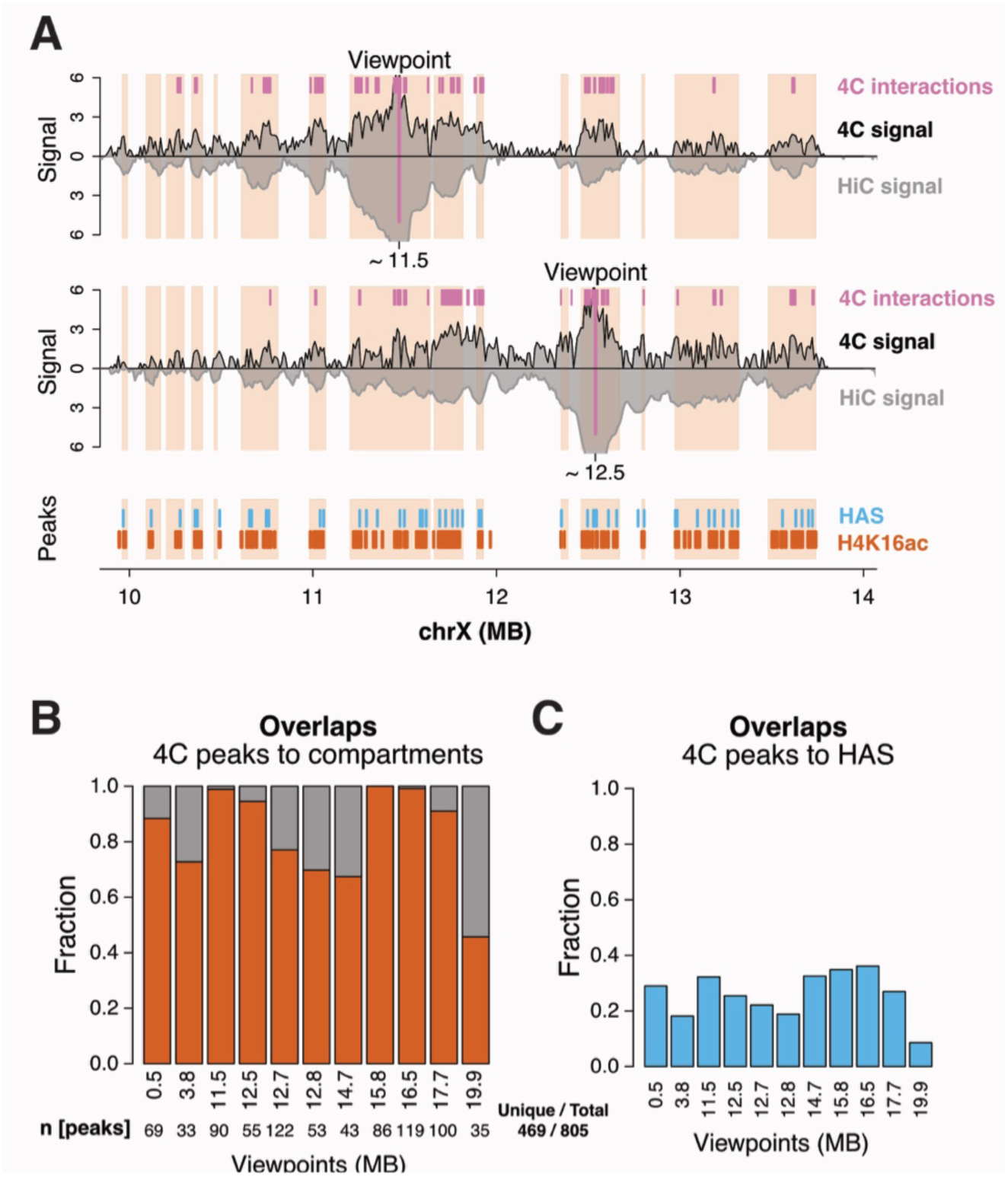
High-resolution 4C profiles reveal 3D interactions between HAS and active domains. **A**) *Top tracks*: 4C (black, S2 cells) and Hi-C (dark grey, male embryos) profiles from viewpoints close to PionX sites at 11.5 (*roX2*) and 12.5 MB (*tomosyn*). 4C interaction peaks are marked by magenta bars and Hi-C active domains by red shaded areas. See Supplementary File 5. *Bottom track*: Peak regions of HAS and H4K16ac are indicated as blue and red bars, respectively. **B**) Fraction of 4C interaction peaks overlapping active (red) and inactive domains (grey) for each viewpoint. Number of peaks is indicated at the bottom. **C**) Fraction of 4C interaction peaks overlapping HAS.

Taken together, the 4C-seq agrees with our Hi-C findings that HAS including PionX sites in active domains mainly interact with other active domains even at megabase distance, thereby ‘jumping over’ intervening inactive compartment domains. This is consistent with a model in which active and inactive domains cluster in the nuclear space and do not intermingle in 3D (Sexton & Cavalli 2015).

### Inducing and disrupting dosage compensation does not alter chromosome conformation

Comparison of Hi-C maps obtained from female and male embryos (this study) as well as from S2 cells upon MSL2 RNAi (Ramírez et al. 2015) suggests that the overall architecture of the X chromosome is independent of sex and dosage compensation. However, given the quantitative nature of the dosage compensation process interactions between individual HAS and adjacent domains may be quantitatively modulated. We therefore extended our high-resolution 4C-seq analysis to monitor differential interactions upon disrupting the complex in S2 cells (by MSL2 RNAi) or inducing DCC assembly in Kc cells (by SXL RNAi). We observed remarkably similar 4C profiles for control and RNAi conditions for both loss and gain of dosage compensation (Figure 4 and Figure 4 – figure supplement 1). Profiles derived from the same viewpoint show very high correlation among biological replicates and among conditions (Pearson´s r ~0.98, r ~0.97, respectively). Hierarchical clustering on the correlation coefficients clearly separates the viewpoints but not the samples from each other (Figure 4 – figure supplement 1C).

**Figure 4.**
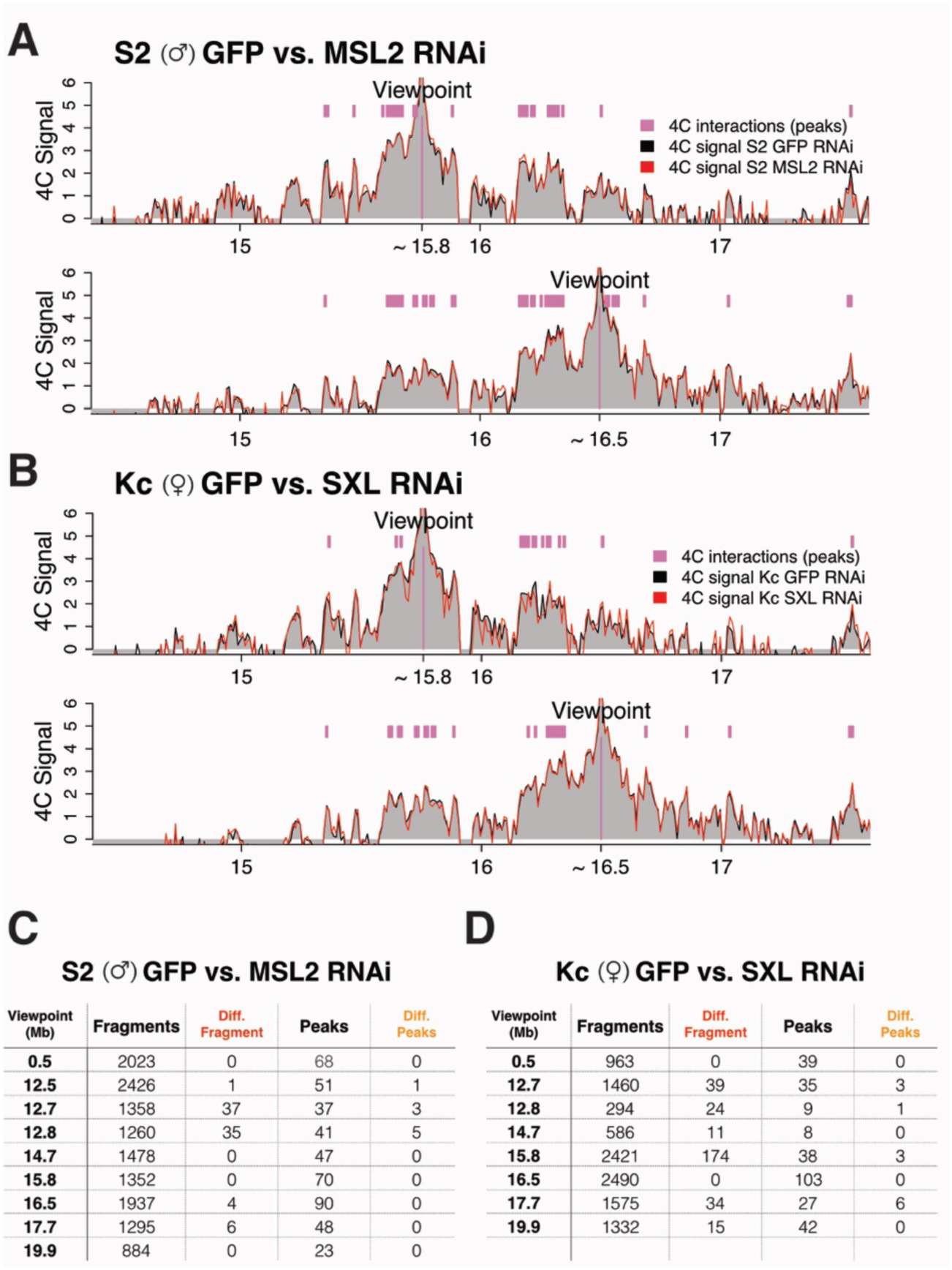
Disrupting and inducing dosage compensation leads to minor changes in 4C signal. **A**) 4C-seq profiles in S2 cells treated with GFP RNAi (black line / grey shading) and MSL2 RNAi (red line) from viewpoints close to HAS at 15.8 and 16.5 MB. Interaction peaks are marked by magenta bars. **B**) Similar graphs as A) but in Kc cells treated with GFP RNAi (black line / grey shading) and SXL RNAi (red line). **C**) Summary statistics (DESeq) of differential 4C fragments between GFP vs. MSL2 RNAi-treated S2 cells and **D**) between GFP vs. SXL RNAi-treated Kc cells. The number of fragments investigated, the differential fragments, the number of interactions (peaks), as well as differential interactions are indicated (cutoff: FDR < 0.01; log2(FC) > 1; n = 2). See Supplementary File 6.

In order to find quantitative differences we analyzed the 4C data by DESeq, as described previously (Ghavi-Helm et al. 2014; Klein et al. 2015)(**Supplementary File 6**). We compared the RNAi samples from each cell line (S2: GFP vs. MSL2 RNAi and Kc: GFP vs. SXL RNAi) to exclude distortions due to the extensive genomic copy number variations between the cell lines. On average, less than one percent of the 4C fragments were significantly different upon depletion of MSL2 in S2 cells (Figure 4C) and for most of the viewpoints the interaction peaks did not change significantly. Two out of the nine viewpoints profiles show a few more differential fragments (i.e. ~3% at 12.7 and 12.8 MB), however these changes are likely explained by local variability of adjacent low count fragments and also by relatively lower replicate correlations (Figure 4 – figure supplement 1C).

Likewise, upon induction of DCC assembly in Kc cells on average only three percent of the contacts changed without clear directionality (Figure 4D and Figure 4 - figure supplement 1B). We note that the number of differential fragments (not peaks) is higher in the Kc cell datasets compared to S2 cells. However, these fragments usually have low read counts compared to the peak regions and quite often adjacent fragments associated with the same gene change direction. Therefore, we suspect that these differences are, although statistically significant, biologically not relevant. In summary, we conclude that neither DCC disruption nor induction alters three-dimensional 4C profiles from individual viewpoints close to PionX signatures, in agreement with previous conclusions from Hi-C analyses.

### The DCC activates genes in close intra- and inter-domain 3D proximity

Our 4C analysis revealed that HAS-associated long-range X-chromosomal interactions occur independently of a functional DCC. As a correlate, the conformation of the X-chromosome may act as a pre-existing scaffold on which the DCC acts. The functional consequences of an individual long-distance contact are difficult to evaluate, given the multitude of interactions of many HAS throughout the active compartment (Figure 2A). Indeed, when dosage compensation is induced in female cells by depletion of the SXL repressor we find that most genes that are activated are less than about 10 kb away from a HAS (Figure 5A). However, under those conditions the newly formed DCC initially binds only a subset of HAS, with PionX signature (Villa et al. 2016)(**Supplementary File 7**). Many domains harbor only a single such induced binding site (15 out of 21; Figure 5 – figure supplement 1A), allowing us to investigate the effect of an isolated binding event on the transcription of surrounding genes. We find that genes located within 50 kb from these sites get significantly higher activated compared to more distal genes (p-value < 10^-7^, two-sided Wilcoxon rank sum test; Figure 5A) even though they reside in same domain with an average size of 200 kb (Figure 5 – figure supplement 1A).

**Figure 5.**
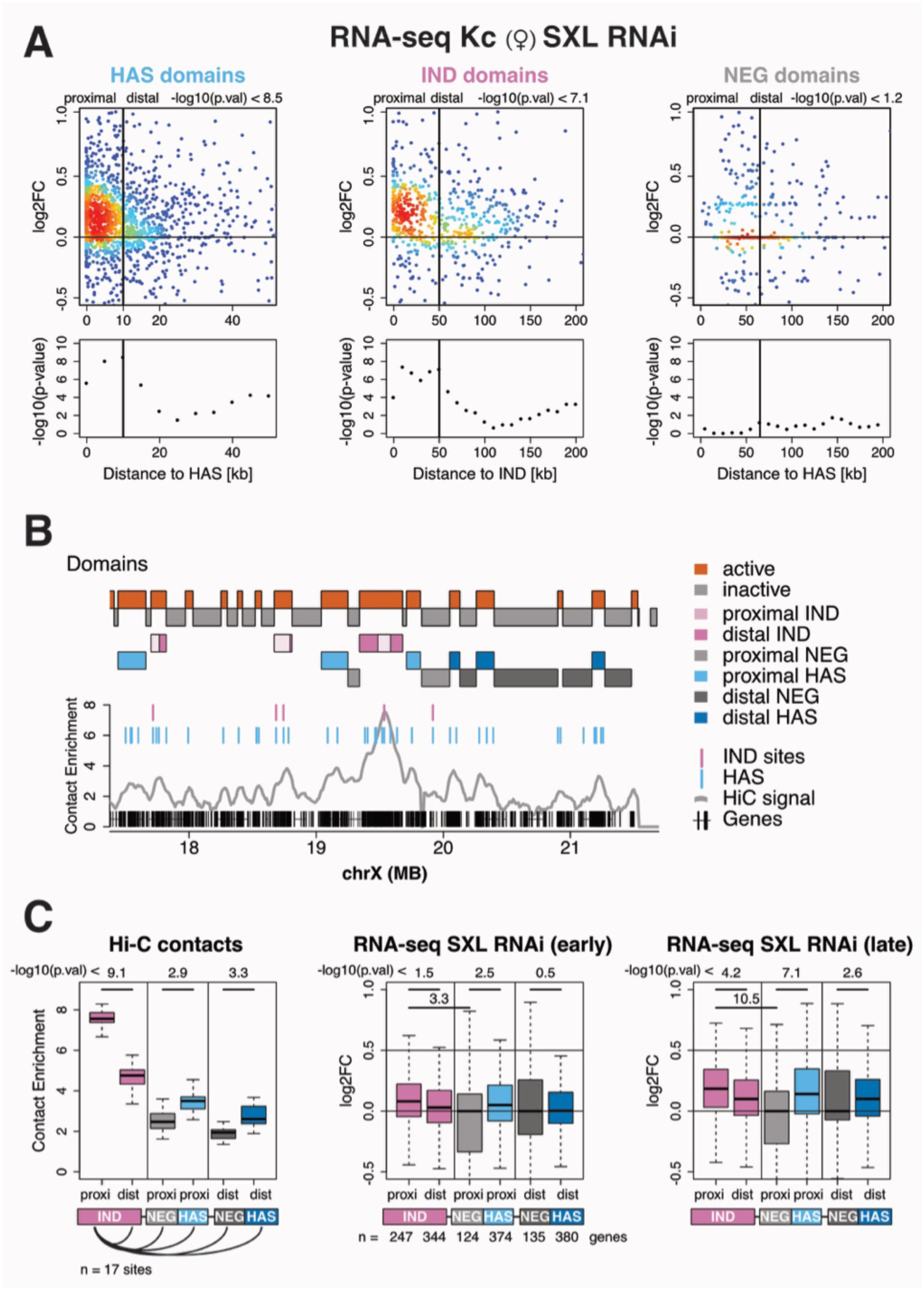
The Dosage Compensation Complex acts locally around High Affinity Sites. **A**) Distance-dependent transcription activation upon SXL RNAi in Kc cells. *Top*: RNA-seq log2 Fold Change (SXL vs. GFP RNAi) related to the distance of genes to HAS (*left* and *right*) and SXL RNAi induced MSL2 binding sites (IND, *middle*) located in domains overlapping HAS (*left*; n=1656), induced sites (*middle*; n=674) or inactive (NEG) compartment (*right*; n=379). Thresholds for proximal and distal genes are indicated as vertical lines and corresponding p-values are calculated by two-sided Wilcoxon rank sum test. *Bottom*: -log10(p-values) [y-axis] justifying the threshold chosen for separating proximal and distal genes [x-axis]. **B**) Example of Hi-C contact enrichment at viewpoint overlapping induced MSL2 binding site close to 19.5 MB. Tracks indicate active (red) and inactive (grey) domains as well as domains proximal (light magenta) and distal (dark magenta) to induced sites (IND), proximal and distal negative control inactive regions (NEG, light and dark grey), proximal and distal HAS regions without induced site (light and dark blue) and average Hi-C contact enrichment (grey profile) from the proximal region as viewpoint. ‘Induced site domains’ larger than 100 kb and other domains larger than 50 kb are considered. **C**) *Left*: Intra-domain and inter-domain Hi-C contact enrichments from domains with induced MSL-2 binding sites. Proximal and distal domains are the same as in B. *Middle and Right*: RNA-seq log2 Fold Change (SXL vs. GFP RNAi) of genes located in the same domains as in B. Early indicates 3-6, whereas late 6-9 days RNAi treatment.

We wished to explore whether activation could also occur through long-range, inter-domain interactions. We therefore scanned the profiles for situations where a domain bearing an induced site was in spatial contact with an active domain that lacks such a site, but separated by inactive compartment. We investigated 14 ‘induced site domains’ (carrying 17 sites) that are larger than 100 kb (see Figure 5B and Figure 5 – figure supplement 1B for examples). Within each domain, a clear distance-dependence in contact enrichment from the induced site viewpoint can be seen, but also contacts ‘jumping over’ the inactive compartment into the neighboring active domain are scored (Figure 5C, **left panel**). The activation of genes within the induced site domain correlated with their distance from the DCC binding site. Remarkably, we found that genes in the neighboring domain were activated to a similar extent as proximal genes within the same induced site domain (Figure 5C, **middle and right panel**). We also distinguish more distal HAS domains that are induced only later during the time course of RNAi (early: 3-6 vs. late: 6-9 days). This suggests that genes in a separated domain are brought into the radius of DCC activation by chromosome folding and that the range of DCC operation should be considered in nuclear space rather than on a linear chromosome map.

## Discussion

### Constitutive transcription organizes chromosome conformation in *Drosophila*

Previous work had shown that the three-dimensional chromosome topology in *Drosophila* cell lines correlates well with chromatin states, RNA polymerase II binding and transcription (Ulianov et al. 2016). It appears that, at least in *Drosophila*, the conformation of chromosomes is an emergent property due to the underlying gene activity pattern. *Drosophila* genes are clustered in constitutively transcribed regions (‘yellow’ chromatin) interspersed by gene-poor, long intron-containing, inactive regions (‘black’ chromatin). In agreement, we find the active compartment marked with H3K36 tri-methylation, H4K16 acetylation and high level of gene expression (RNA-seq). The clustering of ‘alike’ chromatin in space not only leads to the formation of TADs, but is also evident at the level of compartments, which are formed by inter-domain interactions between domains of similar activity status. Indeed, A/B compartments in human cells can be modeled based on DNA methylation, DNase hypersensitive sites or single cell ATAC-seq (Fortin & Hansen 2015).

Our conclusion from the Hi-C data that segments of inactive domains engage in more internal contacts than those in active regions, which appear more extended, agrees with earlier analyses of polytene chromosomes of larval salivary glands. There, the more extended interbands correspond to active, whereas the compact bands reflect the inactive TADs (Ulianov et al. 2016; Eagen et al. 2015).

The fact that the three-dimensional interaction network appears mainly determined by constitutive gene activity explains why we and others (Ramírez et al. 2015) find no major differences in the chromosome conformation of female and male cells and no changes due to dosage compensation. Dosage compensation prominently fine-tunes housekeeping gene transcription (Gilfillan et al. 2006). According to a plausible model the high affinity binding sites for the MSL-DCC evolved from CT-rich intronic splice enhancer precursor sequences (Quinn et al. 2016). Being anchored close to or within active genes the DCC can profit from the hard-wired chromosome compartment organization to contact distant, active chromosomal regions. Basing the viewpoints of our high-resolution 4C-seq study as closely as possible to selected HAS, we zoomed in on the interaction profiles of HAS. We found that relatively short HAS-containing fragments interact widely with active domains, independent of the presence of another HAS in target chromatin. Apparently, the MSL-DCC bound to HAS can come into contact with many loci in the active compartment in their 3D vicinity.

### Extension of active domains in males may be due to DCC-dependent chromosome looping

Although the conformation of the X chromosome in male and female embryos is highly similar, the Hi-C analysis numerically highlighted a shift of interaction frequencies of fragments in the active compartment. Occasionally, this even leads to compartment switching, i.e. cases where homologous chromosomal loci reside in different compartments in male and female nuclei. Extensive switching of entire domains between active and inactive compartments had been described during lineage specification in human cells (Dixon et al. 2015). Since *Drosophila* embryos of different sex mainly consist of the same cell types (except for germ line), one would not expect such differences between female and male Hi-C datasets. The main difference between male and female X chromosomes is that the single X in males is dosage-compensated, which involves enhanced H4K16 acetylation at transcribed gene bodies. H4K16ac also marks promoters of constitutively active genes in both sexes (Feller et al. 2011; Lam et al. 2012), which is why H4K16ac is also enriched in the active compartment in female cells. We documented several cases where a compartment boundary within a transcribed gene was shifted on the male X chromosome such that the gene body was now part of the active compartment. This modulation of chromosomal interactions correlated with high levels of H4K16 acetylation. The broad distribution of H4K16ac may interfere with contacts that define the inactive compartment. Alternatively, an extension of the active compartment may be explained by increased interactions from within the active domain to neighboring chromatin. In a popular scenario the MSL-DCC binds tightly to HAS and then reaches out by chromatin looping to acetylate active chromatin marked by H3K36me3 (Larschan et al. 2007). In such a case the enhanced contacts may reflect the act of acetylation, rather than the presence of H4K16ac itself. The same principle would lead to a globally increased number of contacts within the active compartment on the male X chromosome.

### PionX sites refine X chromosome conformation

One motivation for the current study was to test the hypothesis that PionX sites, which we had identified as first contact points of the DCC on the X chromosome (Villa et al. 2016), modulate the chromosomal conformation in ways that would facilitate dosage compensation. PionX sites do not appear to be necessary for all dosage compensation events since most active domains do not contain such sites. High resolution 4C-seq, which highlights interactions between PionX and adjacent active domains, revealed that the chromosomal contacts of segments harboring PionX motifs are identical in male and female cells and after acute perturbation or induction of dosage compensation. However, averaging across all long-range, inter-domain PionX-PionX contacts we identify slightly increased, more focused interactions in male embryos compared to females, which may point to an additional refinement of folding of the dosage-compensated X chromosome. This observation agrees with earlier DNA FISH studies, where very distal HAS (carrying PionX motif) were closer in 3D in male embryonic nuclei in an MSL2-dependent manner (Grimaud & Becker 2009).

### The MSL-DCC activates target genes by chromosome looping from HAS

Due to the clustering of genes on the X chromosome and the fairly even distribution of HAS, the genes that are subject to dosage compensation are commonly within on average 10 kb of the next HAS. The clear distance-dependence of gene activation (Straub et al. 2008; Ramírez et al. 2015) can now be interpreted in the context of chromosome organization. In order to evaluate the radius of action of an individual HAS we took advantage of our recent observation (Villa et al. 2016) that during early times of ectopic induction of dosage compensation in female cells the DCC preferentially binds to a subset of HAS with PionX signature. Combining domain information and PionX sites we investigated the function of single strong MSL2 binding sites within an active domain. Genes within a 50 kb window around the binding site were activated significantly greater than more distant ones. This determines the functional radius of such a site.

The analysis of gene activation within a domain does not reveal whether the activation over distance involves looping or linear diffusion of the MSL-DCC from a HAS, possibly facilitated by other HAS or secondary elements of lower affinity. However, the observation that primary binding of MSL-DCC to a PionX site in one domain can lead to activation of genes in a second domain that is separated by a large inactive domain strongly argues in favor of a looping mechanism. Such spatial propagation is facilitated by the compartment organization of the chromosome, which is formed by association of active and inactive domains in space (Figure 6). The functional radius of DCC bound to a given site has to be defined as a volume within the active part of the chromosome. The fact that activation can occur over several hundreds of kilobases by spatial looping and yet most genes have a HAS within tens of kilobases suggests a high level of functional redundancy and flexibility in the precise mechanism of activation.

**Figure 6.**
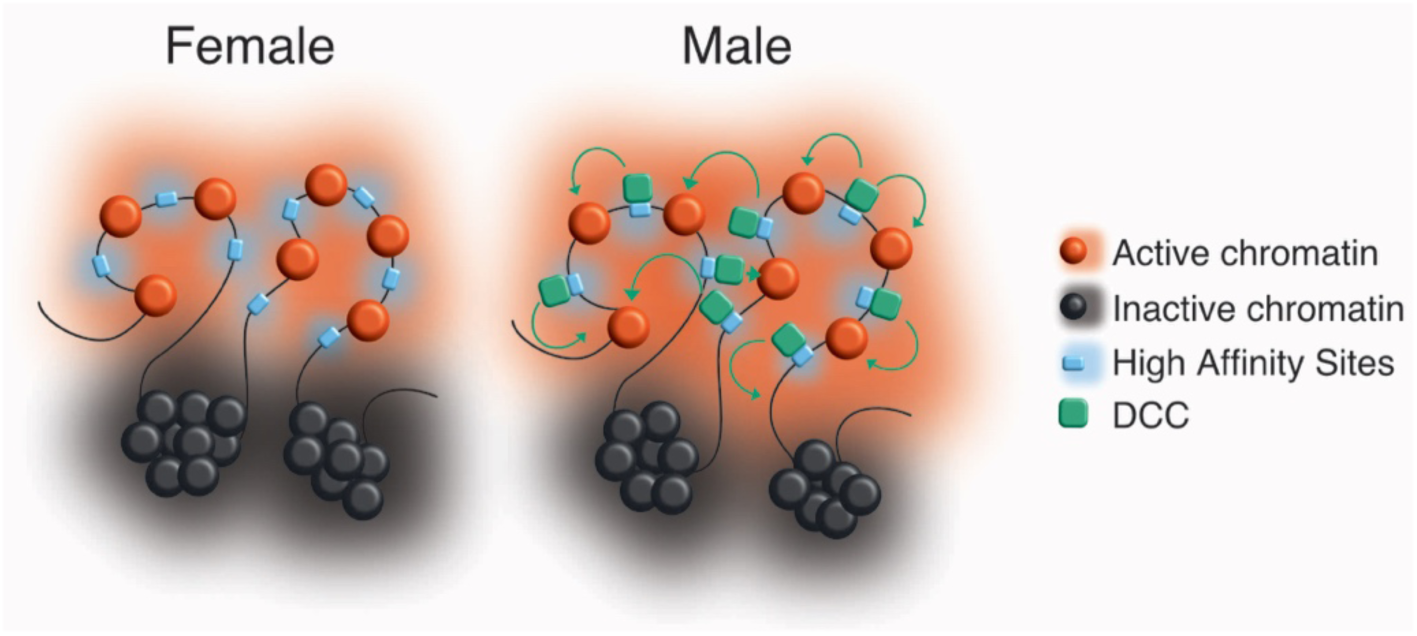
Model of dosage compensation in the context of nuclear architecture. Chromosomes are compartmentalized into active (*red*) and inactive (*black*) domains. This hard-wired architecture is probably determined by general gene activity of constitutive genes independently of sex and dosage compensation. HAS (*blue*) are located in the active compartment allowing for the DCC (*green*) to act on genes in the local spatial proximity (green arrows). DCC assembly only occurs in males (*right*) because of the absence of MSL2 in females (*left*). The process of DCC-mediated acetylation may influence the existing contacts that occasionally leads to extension of the active compartment.

## Conclusions

Analyzing new Hi-C and 4C data we define the compartment structure of the X chromosome in female and male *Drosophila* embryos using principal component analysis. We confirm previous findings of similar nuclear architecture in the sexes obtained in cell culture lines. The chromosome conformation is almost identical in both sexes and not modulated by dosage compensation, a major regulatory principle of male cells. The crucial regulator of dosage compensation, MSL-DCC, profits from a preset chromosome organization. The primary interaction sites for the MSL-DCC (HAS, CES) are evenly distributed within the active compartment, such that they are within a few tens of kilobases from each target gene. Inside an active domain, the strength of transcription activation weakens in a distance-dependent manner. However, the MSL-DCC can take advantage of the compartment organization of *Drosophila* chromosomes and contact active chromatin even if it is a megabase away on the linear chromosome map and separated by inactive domains. These findings establish that the MSL-DCC acts via spatial looping between chromosome segments within the active compartment and that its functional radius must be defined in space, rather than on the linear chromosome map.

## Materials and Methods

### Hi-C analysis

The sex-sorted embryo Hi-C data were generated in biological duplicates and is available from GSE. The embryo sorting was based on a Y chromosomal GFP reporter (y[1]w[67c23]/Dp(1;Y)y[+],P{w[+mC]=ActGFP}SH1) described in (Hayashi 2010). The flies were raised in standard cornmeal yeast extract media at 25°C. Embryos were collected in 0.03% Triton X-100, 0.4% NaCl 16-18 hr after egg laying, then dechorionated for 5 min in fresh bleach. ~3000 GFP^+^ (male) and GFP^-^ (female) embryos each were sorted with a Union Biometra COPAS Large Particle Sorter, and then processed for Hi-C as in (Sexton et al. 2012).

Paired-end reads were mapped separately to the reference genome (BDGP5.74) using bowtie2 (2.0.2) with “-local” settings (**Supplementary File 1**). In general, all Hi-C data processing and analysis steps were carried out using the Homer software package (Heinz et al. 2010). In detail, paired-end tag directories were generated either by pooling technical (sequencing) replicates for biological replicate comparisons or also pooling biological replicates to maximize coverage for each condition. Reads were extensively filtered with the recommended parameters, i.e. “-tbp 1 -removePEbg -fragLength 500 -restrictionSite GATC -both - removeSelfLigation -removeSpikes 10000 5” (**Supplementary File 1**). Background models, coverage-normalized (“-simpleNorm”) as well as coverage- and distance-normalized correlation matrices (“-corr”) were created using various resolutions (“-res”) and smoothing (“-superRes”) windows, from which 10 kb resolution with 50 kb smoothing was chosen for downstream analysis. Contact enrichment values were always extracted from the coverage-normalized matrix. The genome was divided into two compartments based on principal component analysis of the correlation matrix (“runHiCpca.pl”), where H4K16ac regions (peaks) served as a seed to define the sign of the PC1 values (**Supplementary File 2**). Active or inactive domains were defined as continuous regions with positive or negative PC1 values. To directly compare female and male Hi-C profiles, correlation between interaction-profiles of individual loci (fragments) were calculated among the samples (“getHiCcorrDiff.pl”).

Downstream analysis (overlaps, densities, statistical tests etc.) and plotting (heatmaps, profile plots, boxplots etc.) was performed using R and Bioconductor packages. To test whether there is a difference in Hi-C contact enrichment values between compartments or subsets of domains non-parametric, two-sided Wilcoxon rank sum tests were performed.

### 4C-seq experimental procedures

4C-seq was carried out on S2-DRSC cells (DGRC stock: 181) and Kc167 (DGRC stock: 1) in biological duplicates. RNAi was performed as described previously (Straub et al. 2005) using the following primers for dsRNA PCR template generation:

GFP-forward: TTAATACGACTCACTATAGGGTGCTCAGGTAGTGGTTGTCG,

GFP-reverse: TTAATACGACTCACTATAGGGCCTGAAGTTCATCTGCACCA;

MSL2-forward: TTAATACGACTCACTATAGGGAGAATGGCCCAGACGGCATAC,

MSL2-reverse: TTAATACGACTCACTATAGGGAGACAGCGATGTGGGCATGTC

SXL-forward: TAATACGACTCACTATAGGGAGACCCTATTCAGAGCCATTGGA,

SXL-reverse: TAATACGACTCACTATAGGGAGAGTTATGGTACGCGGCAGATT

(SXL from DRSC28896; (Alekseyenko et al. 2012)).

After 7 days of RNA interference cells were resuspended in fresh medium and fixed with 1% formaldehyde for 10 min at room temperature. Fixing was quenched by adding glycine (final concentration 125 mM) and by cooling on ice. Cells were collected in a cooled centrifuge, snap-frozen in liquid nitrogen and stored at -80°C. 4C-seq templates were generated on ~60 million fixed cells per replicate using 4-cutter restriction enzymes DpnII and NlaIII, as described previously (Ghavi-Helm et al. 2014). Libraries were amplified using 100 ng 4C-seq templates and Pfu Turbo DNA polymerase in 8 PCR replicates, which were pooled for later analysis. Primers were designed to allow a multiplexing of 48 samples (for 12 viewpoints and 4 conditions) on a sequencing lane (**Supplementary File 3**). Biological replicates were sequenced on two separate lanes of an Illumina NextSeq 500 sequencer. Datasets are available at GSE….

### 4C-seq analysis

4C-seq analysis was essentially based on the FourCSeq Bioconductor package (Klein et al. 2015). In detail, FASTQ files were de-multiplexed using a python script available in the package and aligned to the reference genome (BDGP 5.74) using bowtie2 (2.0.2) with “-local” settings. Reads were filtered for mapping quality (mapq > 1) and non-digested, self-ligated fragments were removed (**Supplementary File 4**). Reads were initially counted for each valid DpnII fragment and obtained read counts per fragment were summed over larger windows (from 2.5 kb up to 10 kb). Z-Scores were calculated at each of these of windows with various minimum count thresholds (“getZScores()”). To identify peaks (significant interactions) a Z-score threshold of 2 and an FDR threshold of 0.05 were applied using the “addPeaks()” function. Final settings of window size and minimum counts were chosen based on the number of peaks obtained (**Supplementary File 5**). Z-scores and peak coordinates were used for plotting 4C profiles and for overlap analysis. To find significant differences among conditions DESeq was run on the counts for each fragment (or window) using the “getAllResults()” function of the FourCSeq package (**Supplementary File 6**). Cutoffs were defined with an adjusted p-value < 0.01 and a change log2(FC)l > 1.

### ChIP-seq datasets and analysis

Datasets related to High Affinity Sites (HAS), CXC-dependent (PionX) and SXL-induced site definitions were described in (Straub et al. 2013) and (Villa et al. 2016). Briefly, HAS were identified as the co-localization of MSL-2 and MLE in *in vivo* ChIP-seq (Straub et al. 2013). PionX sites are *in vitro* DIP-seq MSL-2 sites that lose binding upon deletion of its CXC domain and SXL-induced sites were obtained by hierarchical clustering of MSL-2 sites upon SXL RNAi time-course in Kc cells ((Villa et al. 2016), **Supplementary File 7**).

In parallel to the SXL RNAi experiments in Kc cells, we generated a new set of MSL2 (4 replicates) and H4K16ac (1 replicate) ChIP-seq profiles in S2 cells. These ChIP experiments were performed according to (Schauer et al. 2013) adapted to cell lines (Villa et al. 2016). Briefly, cells were resuspended in ice-cold homogenization buffer and fixed with 1% formaldehyde for 10 minutes at room temperature. After quenching with 125 mM glycine the cells were collected, washed and sonicated in RIPA buffer with a Covaris sonifier (PIP: 140, DF 20%, CB: 200) for 30 min. MSL2 antibody was previously described in (Villa et al. 2012) and H4K16ac was acquired from Active Motif (#39167).

H3K36me3 ChIP-seq was performed on MNase digested chromatin as described previously (Straub et al. 2008) and (Kasinathan et al. 2013) with some modifications. Cells were fixed in 1% formaldehyde for 1 min at 26°C, followed by quenching with 125 mM glycine and washing with PBS. Nuclei were released by resuspending in TM2+ with NP-40. MNase digestion was performed in TM2+IC using 4U MNase (Sigma Aldrich, resuspended in EX50 (Bonte & Becker 1999)) in the presence of CaCl2 for 13 min at 37 °C. Reaction was stopped with EGTA and Triton-X-100, SDS, NaDOC and NaCl was added to final concentration as in RIPA Buffer. MNase digested chromatin was incubated for 1 h at 4°C while slight agitation and chromatin was solubilized by passing ten times through 27 G needle and centrifuged for 30 min at 15,000 g at 4°C. ChIP was performed from soluble chromatin using H3K36me3 antibody (ab9050, Abcam).

50 bp single reads were obtained on an Illumina HiSeq sequencer. Reads were mapped to the reference genome (BDGP 5.74) using bowtie2 (2.0.2) with default settings. Peaks were called over corresponding inputs using the Homer Software Package with the parameters “style factor” for MSL2 and “-style histone -F 2.5” for H4K16ac. The final set of MSL2 peaks were based on peak calling in at least two out of the four replicates. The single replicate of the H4K16ac regions was compared to previously published profile using the same peak finding software with an overlap of ~90% on the X chromosome (Straub et al. 2013).

For the comparison of Hi-C PC1 values to H3K36me3 or H4K16ac reads were counted at each 10 kb genomic fragment and were normalized to the total number of reads as well to input and converted to standardized log2 unit, finally averaged over two biological replicates. Otherwise, input- and coverage-normalized tracks for H4K16ac were generated by the Homer Software Package.

### RNA-seq

RNA-seq was performed on S2-DRSC and Kc167 cells at least in biological duplicates. In RNAi experiments the following primers were used for dsRNA PCR template generation:

GFP, MSL2 and SXL (DRSC28896): see under “4C-seq experimental procedures”; and

SXL from DRSC21490 (Alekseyenko et al. 2012):

SXL-forward: TAATACGACTCACTATAGGGAGAGATCACAGCCGCTGTCC,

SXL-reverse: TAATACGACTCACTATAGGGAGATACCGAATTAAGAGCAAATAATAA

Total RNA was prepared from 4 million cells using RNeasy mini columns (Qiagen) and 2.5 μg of RNA was used for polyA selection. Libraries were prepared using New England Biolabs reagents according to protocols (E7490, E6150, E7525, E6111, E7442, E7335 and (Peleg et al. 2016)). 50 bp single reads were obtained on an Illumina HiSeq sequencer. Alignment was performed using Tophat2 (2.0.5) against the reference genome (BDGP 5.74). Reads were counted at exons and summed for each gene (reference: BDGP5.74.gtf) using the “summarizeOverlaps” function in the Genomic-Alignments package. Counts were normalized using the “estimateSizeFactors()” function in the DESeq2 package. Gene expression levels were calculated as log2(normalized counts + 1) for all the genes, including genes with zero counts. To estimate genes expression changes between RNAi conditions the difference of log2(normalized counts + 1) was taken. To test whether there is a difference in gene expression or response to RNAi between subsets of genes non-parametric, two-sided Wilcoxon rank sum tests were performed.

## Acknowledgements

This work was supported by the European Research Council under the European Union's Seventh Framework Programme (FP7/2007-2013) / ERC grant agreement n°293948. We thank S. Krause for technical support. We thank T. Straub for helpful discussions and S. Krebs and H. Blum (LAFUGA at Gene Center, LMU, Munich) for outstanding sequencing service.

### Competing interest

The authors declare that no competing interests exist.

---

**Figure 1 – figure supplement 1.**
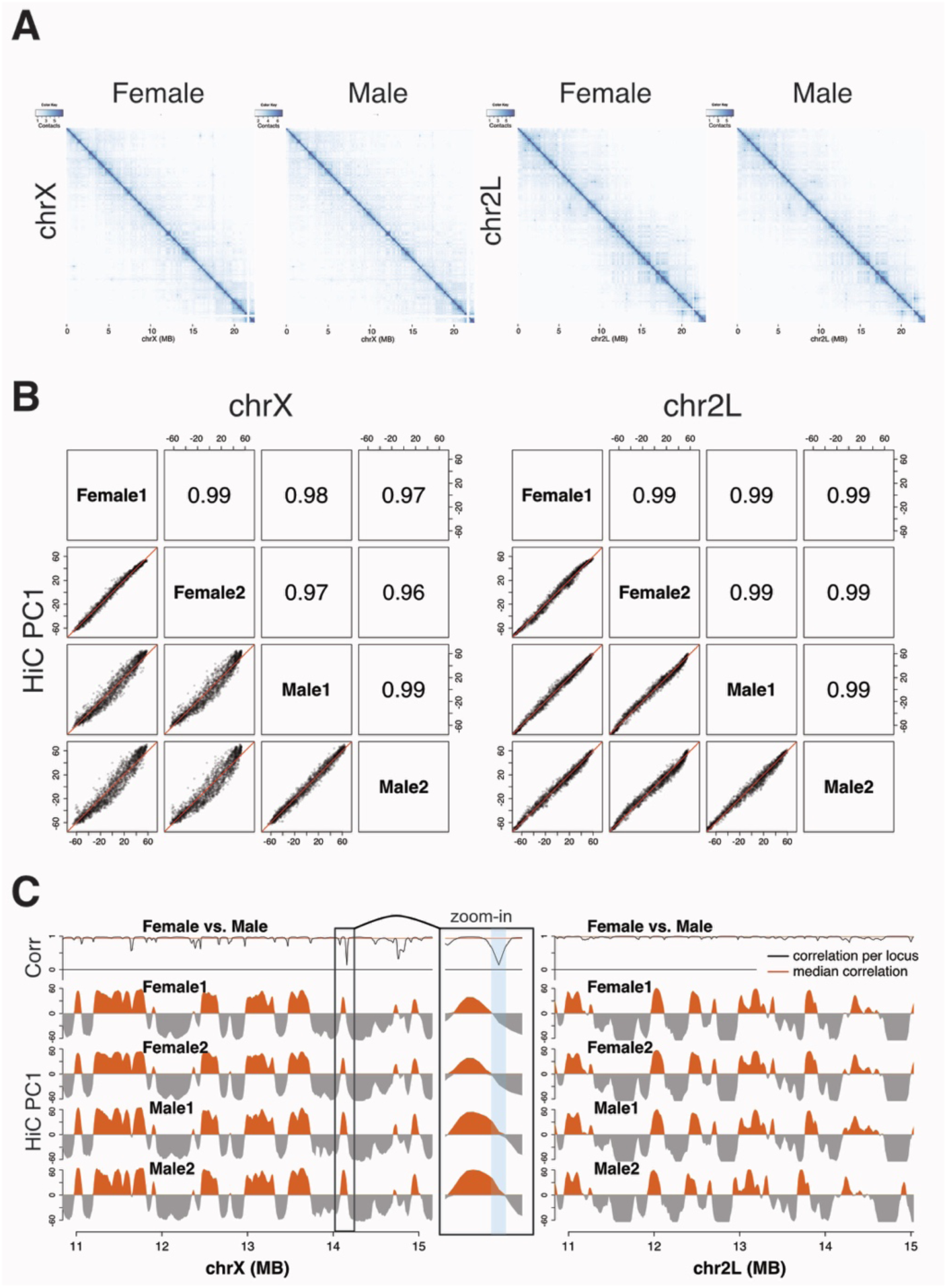
Hi-C contact maps and the first principal component (PC1) of chromosome X and 2L. **A**) Coverage-normalized Hi-C contact maps of chromosomes X and 2L in female and male embryos. **B**) Scatterplots of Hi-C PC1 values for the X chromosome (*left*) and chromosome 2L (*right*) in female and male embryos (2 biological replicates). Pearson´s correlation coefficients are shown in the corresponding upper panels. **C**) Comparison of Hi-C PC1 values for the X chromosome (*left*) and 2L chromosome (*right*) at the region 16-20 MB in female and male embryos (2 biological replicates). Top track shows the correlation of contact enrichment between female and male samples for each locus (fragment). ‘Zoom-in’ shows a view of a locus switching compartments and having low correlation between female and male.

**Figure 1 – figure supplement 2.**
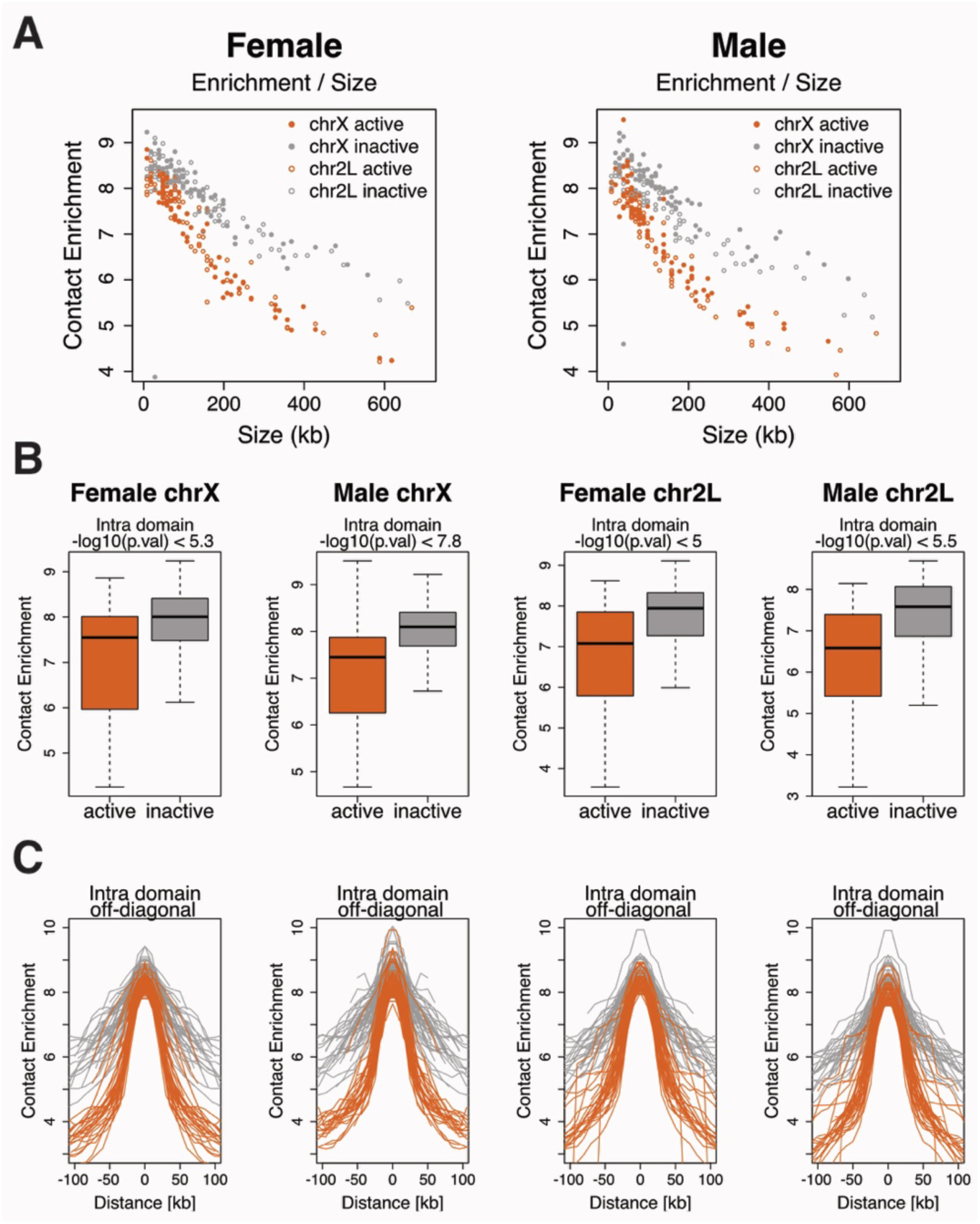
Relationship between domain size and contact enrichment. **A**) Relationship between domain size and average intra-domain contact enrichment for domains located in the active (red) or in the inactive (grey) compartment on chromosome X (solid dots) and 2L (open dots) in females (*left*) and in males (*right*). **B**) Average intra-domain contact enrichments for domains located in the active (red) or in the inactive (grey) compartment. Sample and chromosome names are indicated on the top, p-values are calculated by two-sided Wilcoxon rank sum test. **C**) Off-diagonal intra-domain contact enrichment for each domain located in the active (red) or in the inactive (grey) compartment. Sample and chromosome names are the same as in B). Enrichment values are aligned to the middle of the domain.

**Figure 1 – figure supplement 3.**
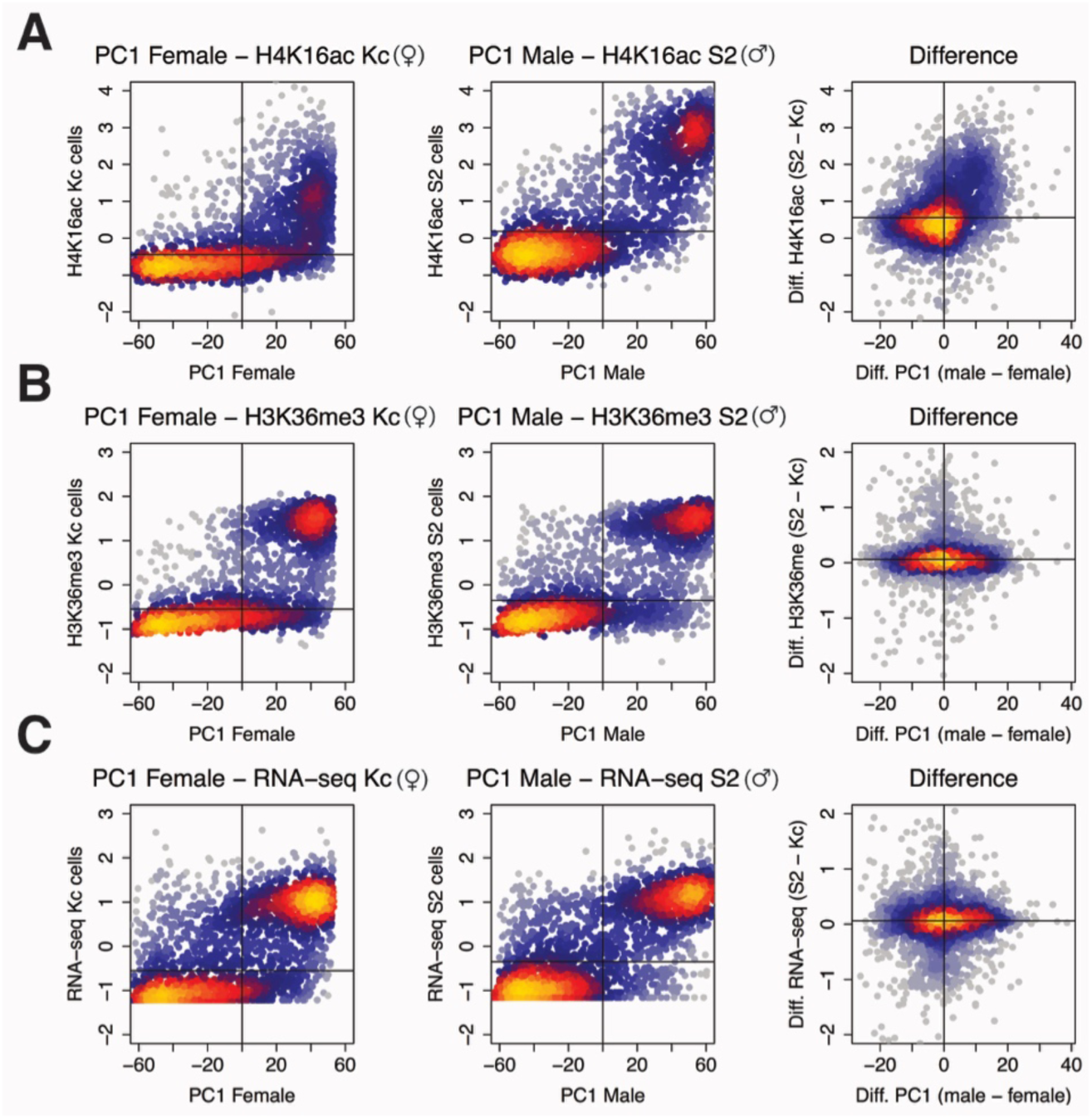
Hi-C fragments with higher PC1 values carry higher H4K16ac levels in males compared to females. **A**) Scatter plot showing the relationship between PC1 (females or males) and H4K16ac (Kc or S2 cells) for individual loci (resolution: 10 kb). **B**) Scatter plot showing the relationship between PC1 (females or males) and H3K36me3 (Kc or S2 cells) for individual loci (resolution: 10 kb). **C**) Scatter plot showing the relationship between PC1 (females or males) and RNA-seq (Kc or S2 cells) for individual loci (resolution: 10 kb).

**Figure 1 – figure supplement 4.**
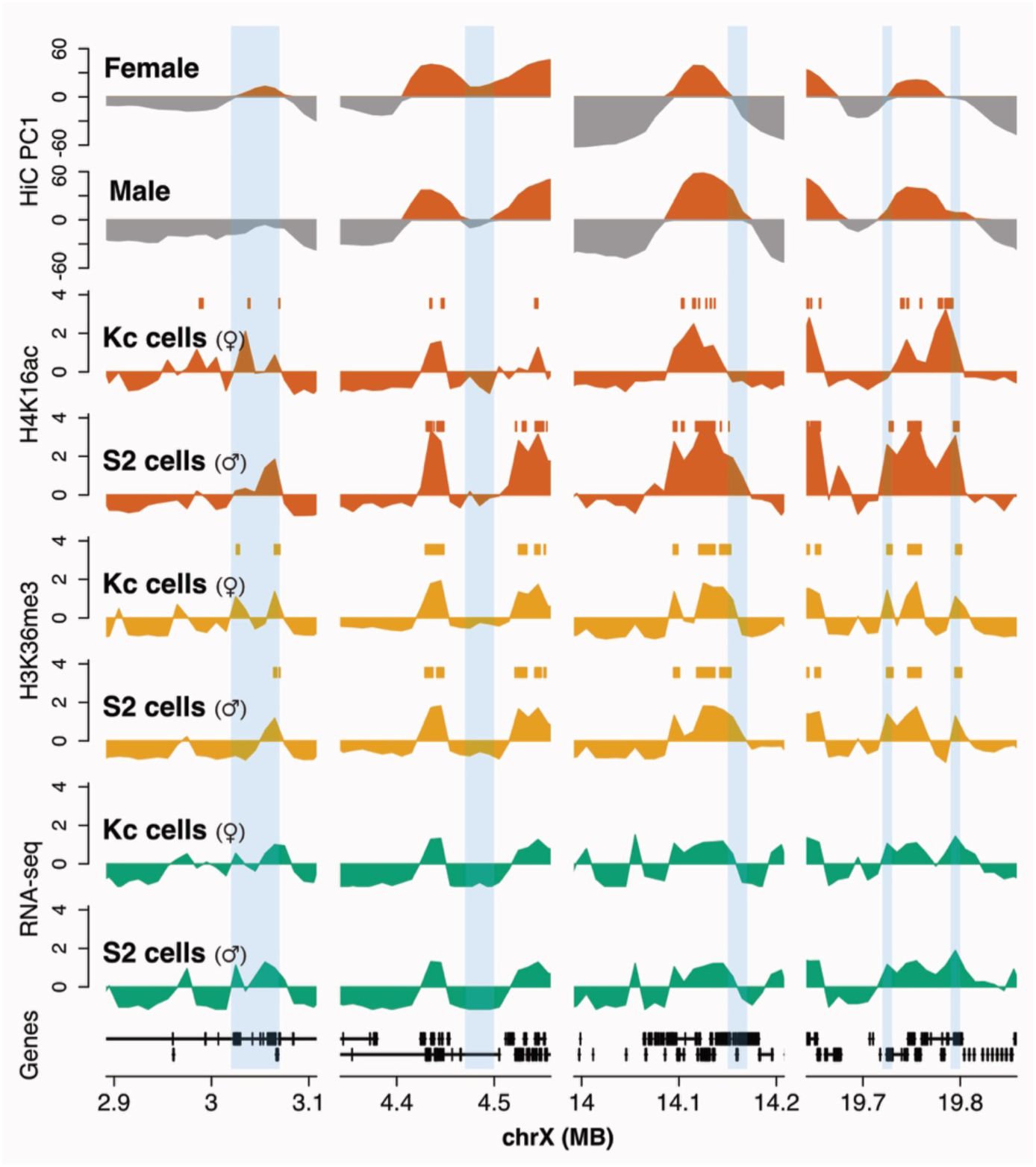
Example regions switching compartments. Tracks indicate PC1 in females and males, smoothed and standardized H4K16ac, H3K36me3 and RNA-seq signal in Kc and S2 cells as indicated. Shaded areas are regions that change the sign of PC1.

**Figure 2 – figure supplement 1.**
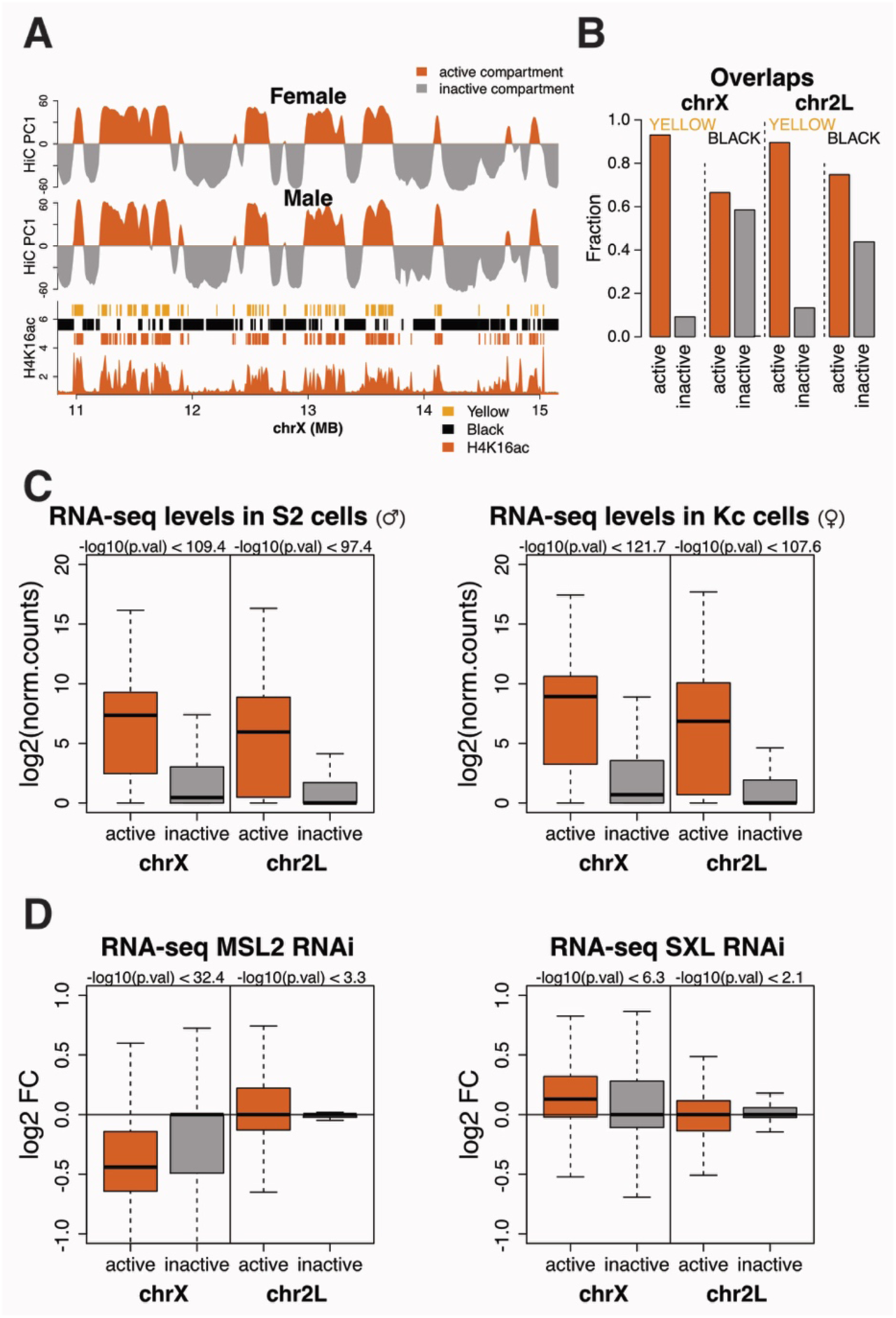
Gene activity in nuclear compartments. **A**) Example region of Hi-C compartment structure and epigenetic domains defined as “yellow” (constitutively active) and “black” (inactive) chromatin (Filion et al. 2010). **B**) Fraction of ‘yellow’ and ‘black’ chromatin domains overlapping with active (red) or inactive (grey) compartment on the X and 2L chromosomes, respectively. **C**) RNA-seq levels [log2(norm. counts)] for genes in the active (red) or in the inactive (grey) compartment on chromosomes X and 2L in S2 cells (left) and in Kc cells (right). P-values are calculated by two-sided Wilcoxon rank sum test (biological replicates n = 2). **D**) RNA-seq log2 Fold Change for the same gene classes as in A) upon MSL2 RNAi in S2 cells (left) and SXL RNAi in Kc cells (right). P-values are calculated by two-sided Wilcoxon rank sum test (biological replicates n = 2).

**Figure 2 – figure supplement 2.**
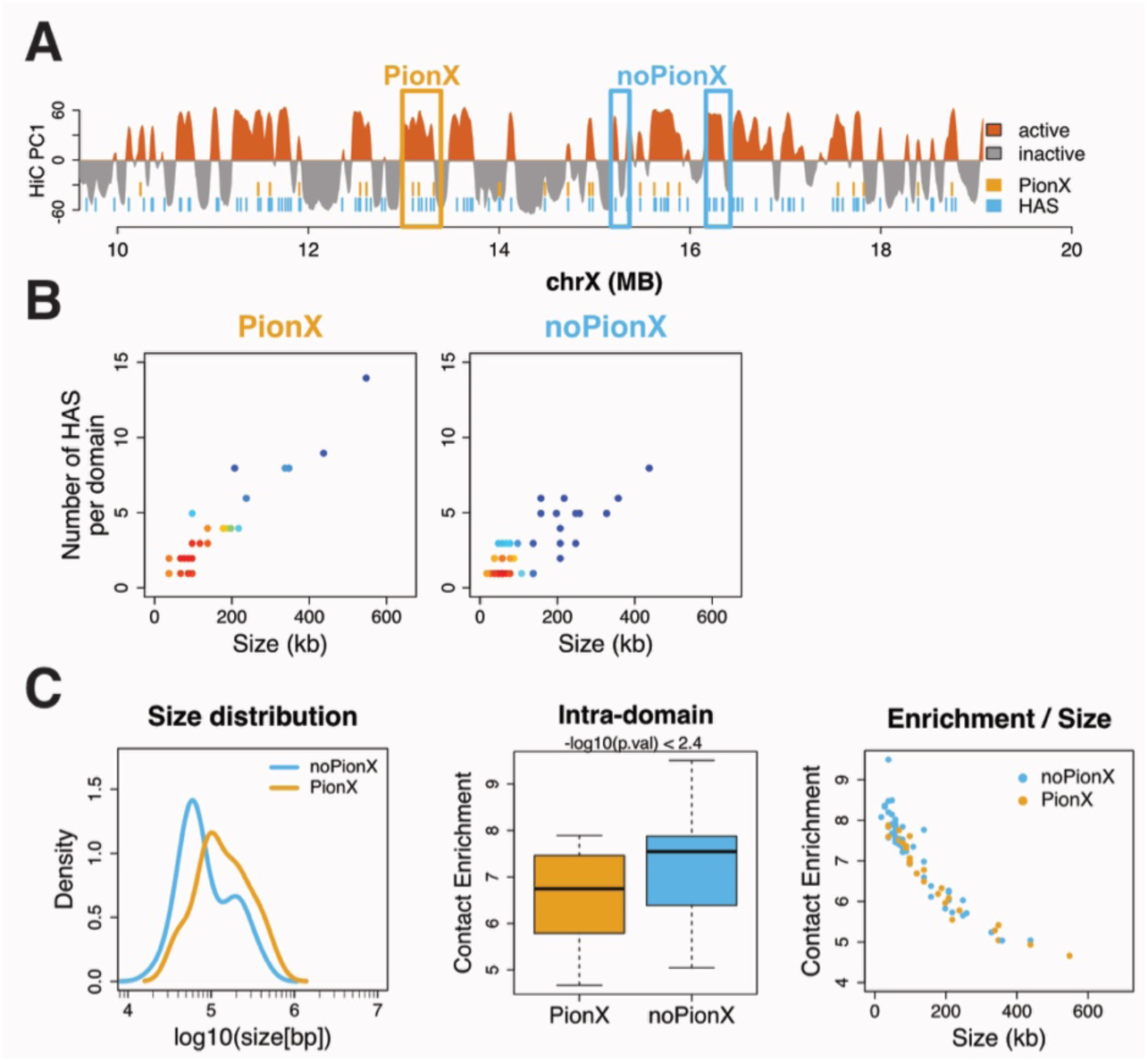
Domains overlapping HAS sub-classes. **A**) Zoom-out view of domains overlapping HAS with (ocher) or without (blue) PionX sites. Example domains are framed. PionX sites and HAS are marked by ocher and blue bars, respectively. **B**) Relationship between the domain size (kb) and the number of HAS in a domain for domains overlapping HAS with PionX (*left*) and HAS without PionX sites (*right*). Color-key: red indicates higher density. **C**) *Left*: Size distribution of domains overlapping HAS with PionX (ocher) and HAS without PionX sites (blue). *Middle*: Average intra-domain contact enrichments for domains overlapping HAS with PionX (ocher) and HAS without PionX sites (blue). P-values are calculated by two-sided Wilcoxon rank sum test. *Right*: Relationship between the domain size and intra-domain contact enrichment for domains overlapping HAS with PionX (ocher) and HAS without PionX sites (blue).

**Figure 2 – figure supplement 3.**
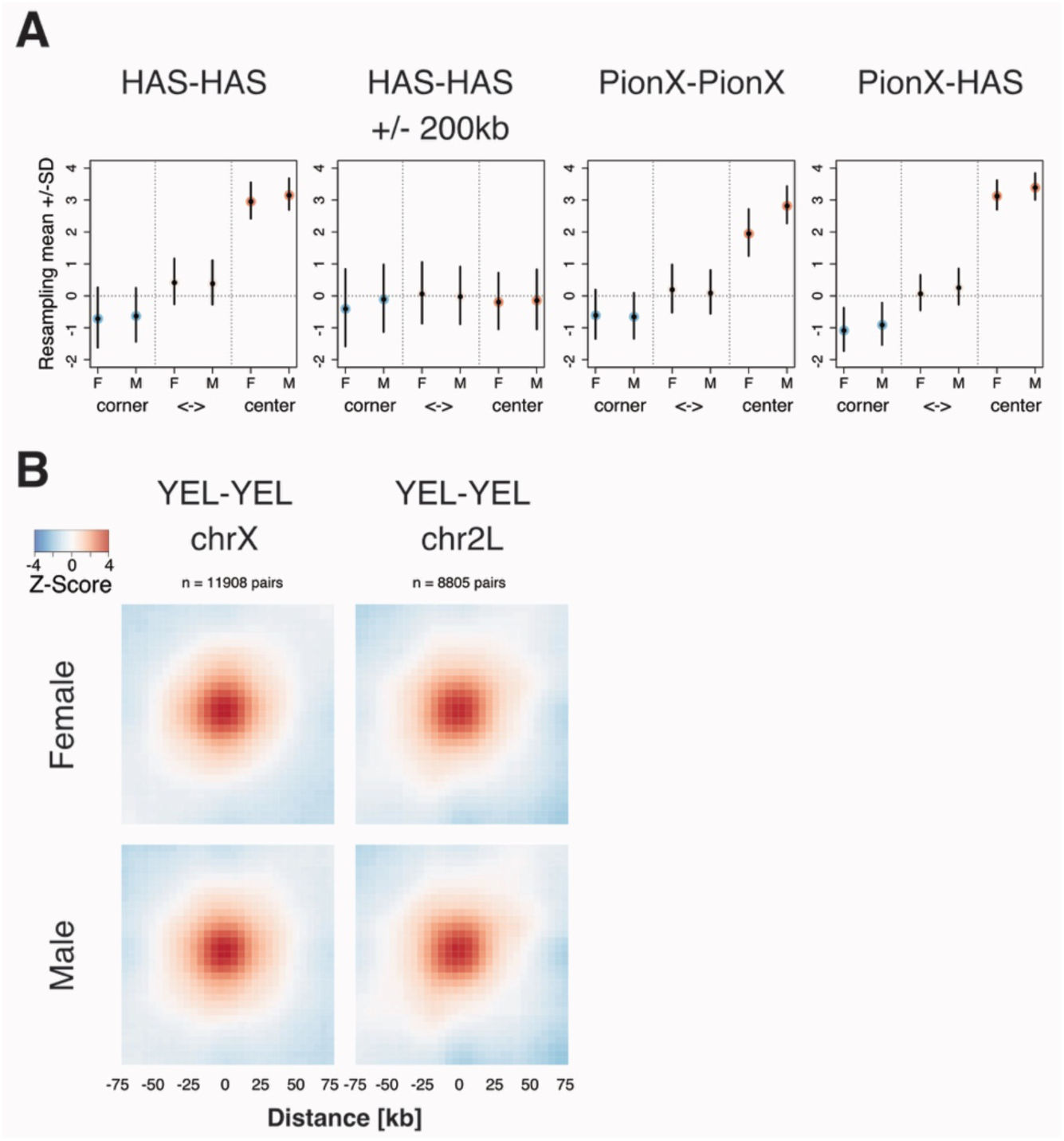
Inter-domain contact enrichment at HAS sub-classes. **A**) Inter-domain average contact enrichment at the corner and center of sub-matrices aligned around pairs of sites (same as in Figure 2D). Mean enrichment and standard deviation derived from iteratively resampled sites are shown (30 sites, 1000 iteration). **B**) Inter-domain average contact enrichment sub-matrices centered around pairs of ‘yellow’ chromatin domains on chromosomes X (left) and 2L (right) (Filion et al. 2010). Interactions between 250 kB and 2.5 MB distance are considered.

**Figure 3 – figure supplement 1.**
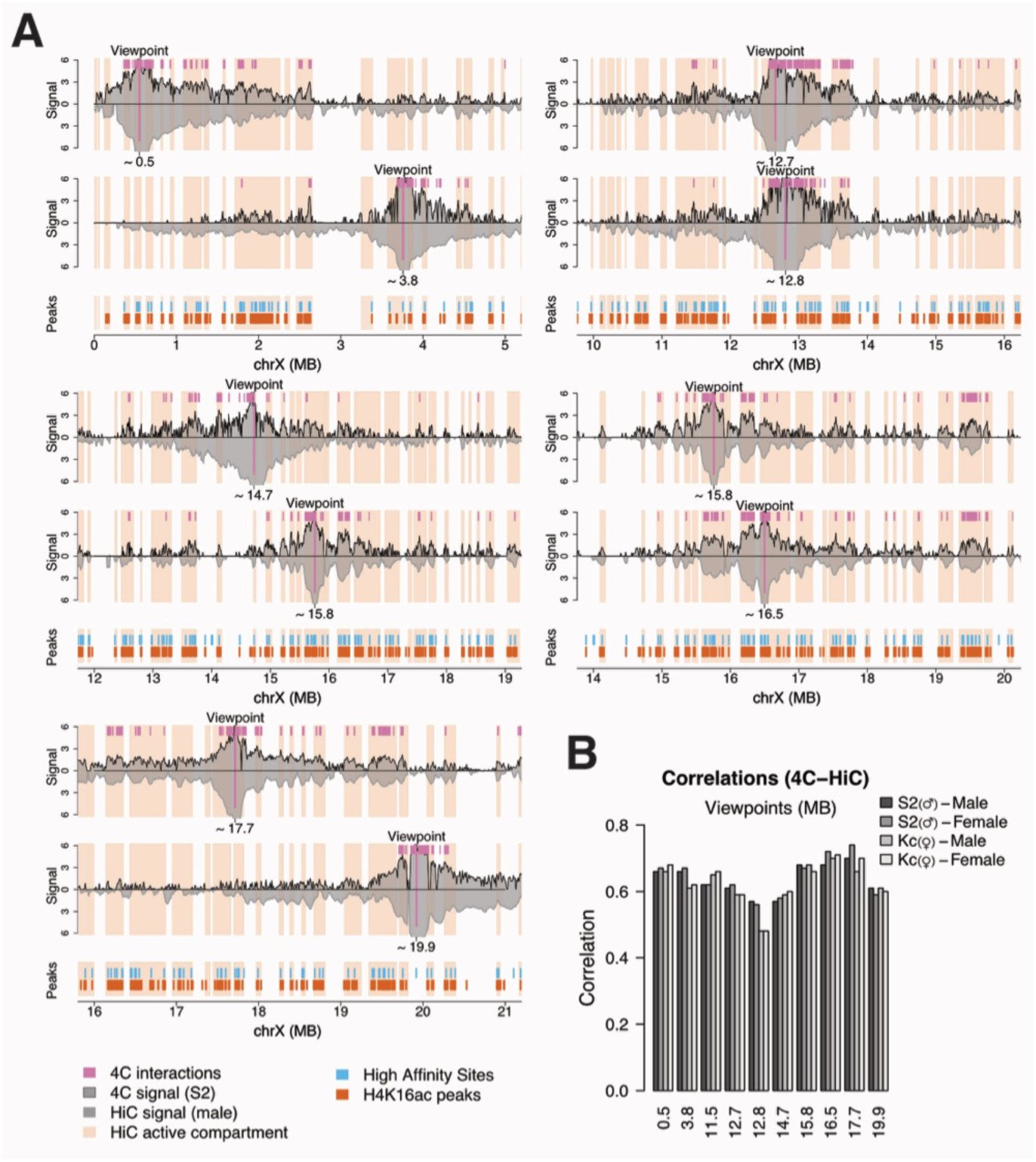
4C profiles at various viewpoints. **A**) Similar plot as Figure 3A for viewpoints close to PionX sites at 14.7, 17.7 and 19.9 MB, close to SXL RNAi induced MSL2 sites at 0.5, 12.7, 12.8, 15.8 and 16.5 as well as close to ‘regular’ HAS at 3.8 MB. **B**) Pearson correlation coefficients for comparison of 4C (S2 or Kc cells) to Hi-C profiles (female or male embryos) at 4C viewpoints.

**Figure 3 – figure supplement 2.**
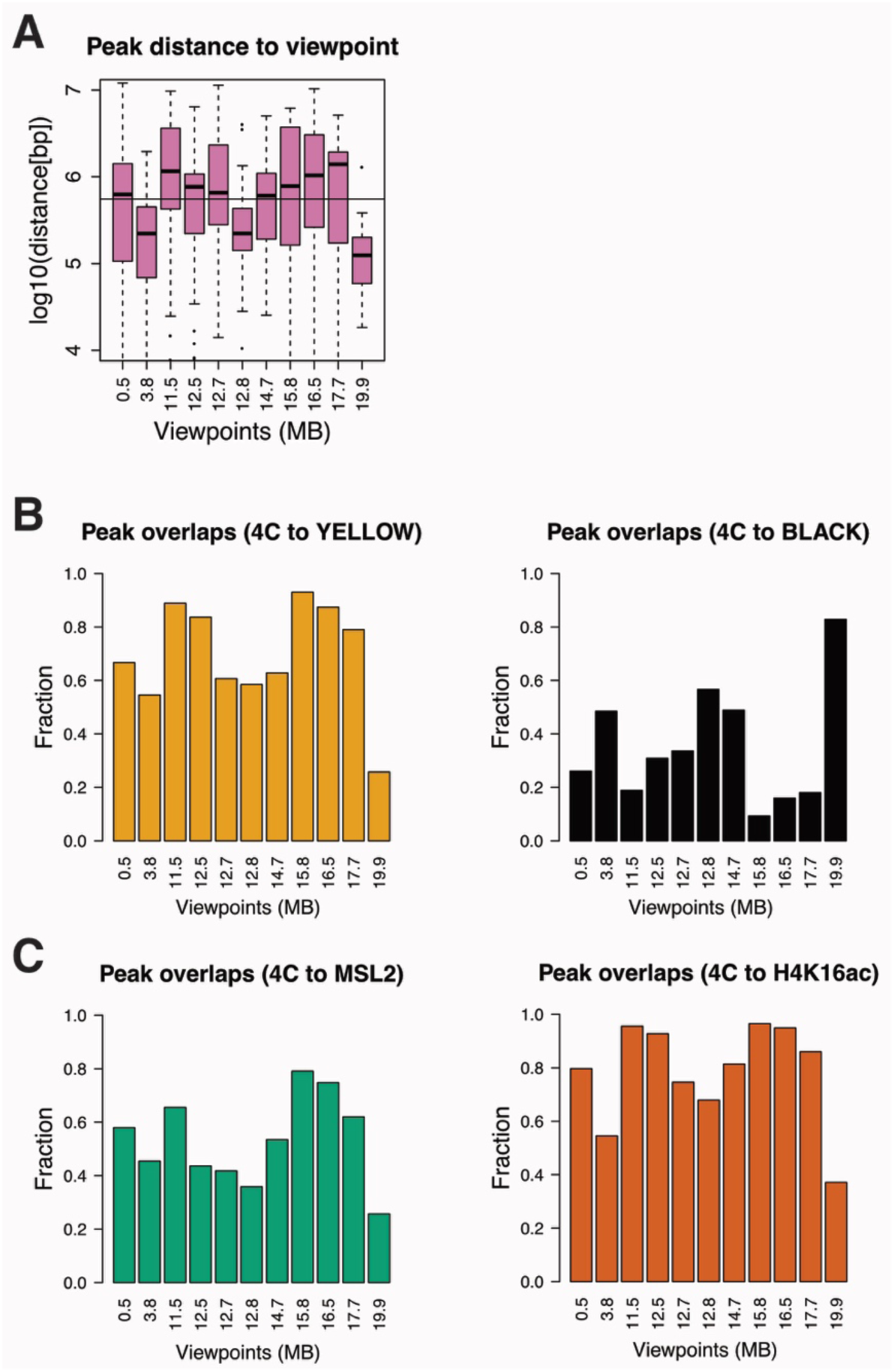
Overlap of 4C interaction peaks with chromatin features. **A**) Distribution of distance (bp) between 4C viewpoint and 4C interaction peaks for each viewpoint. **B**) Fraction of 4C interaction peaks overlapping ‘yellow’ (*left*) and ‘black’ (*right*) chromatin domains for each viewpoint (Filion et al. 2010). **C**) Same as B) for MSL2 peaks (*left*) and H4K16ac (*right*).

**Figure 4 – figure supplement 1.**
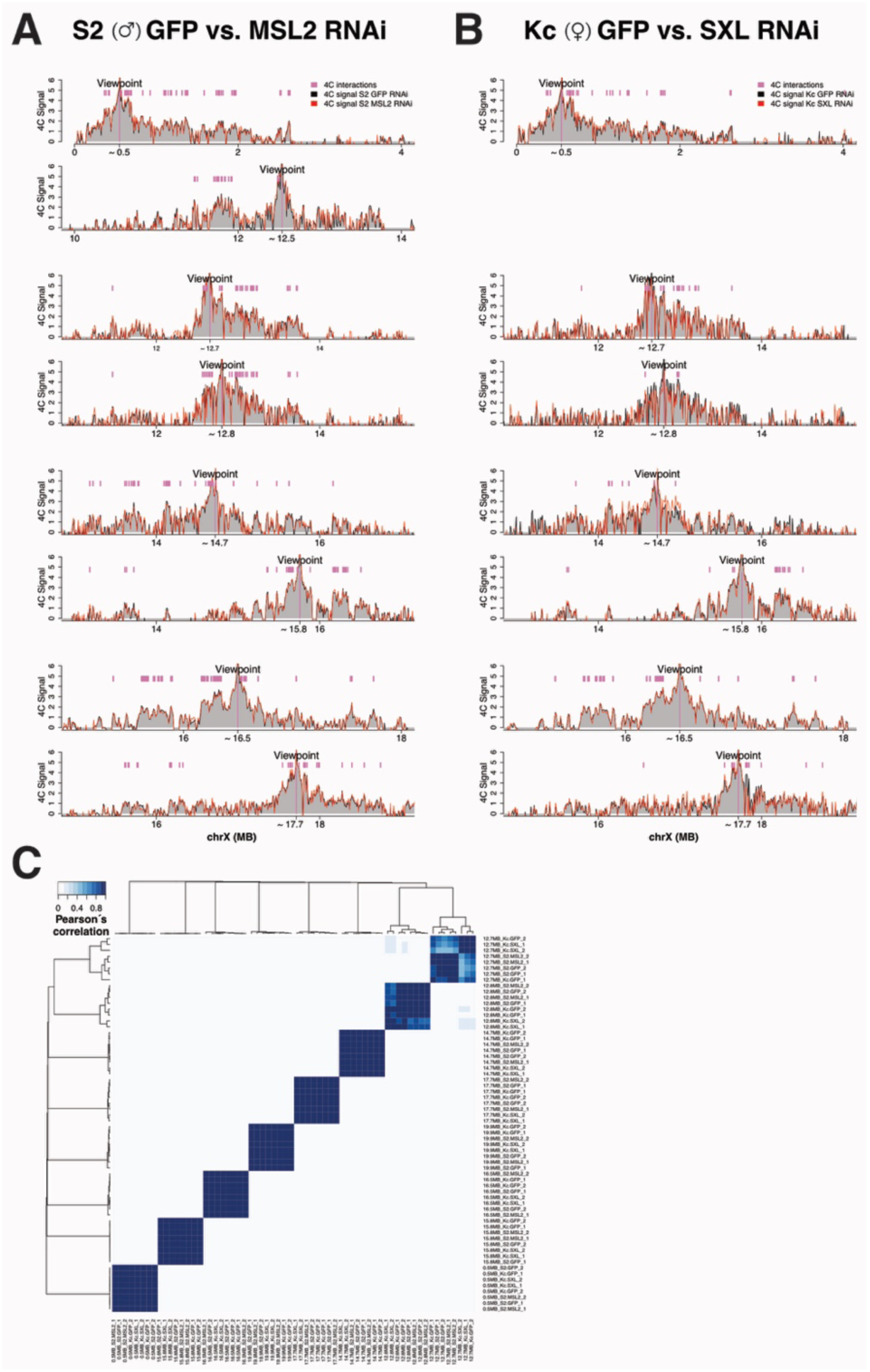
Additional 4C profiles on disrupting and inducing DCC. **A**) and **B**) Similar plots as Figure 4A, respectively for viewpoints close to PionX at 12.5, 14.7 and 17.7 MB and close to SXL RNAi induced MSL2 sites at 0.5, 12.7, 12.8, 15.8 and 16.5 MB. **C**) Heatmap of the Pearson´s correlation matrix for 4C profiles including all the conditions and biological replicates (n = 2).

**Figure 5 – figure supplement 1.**
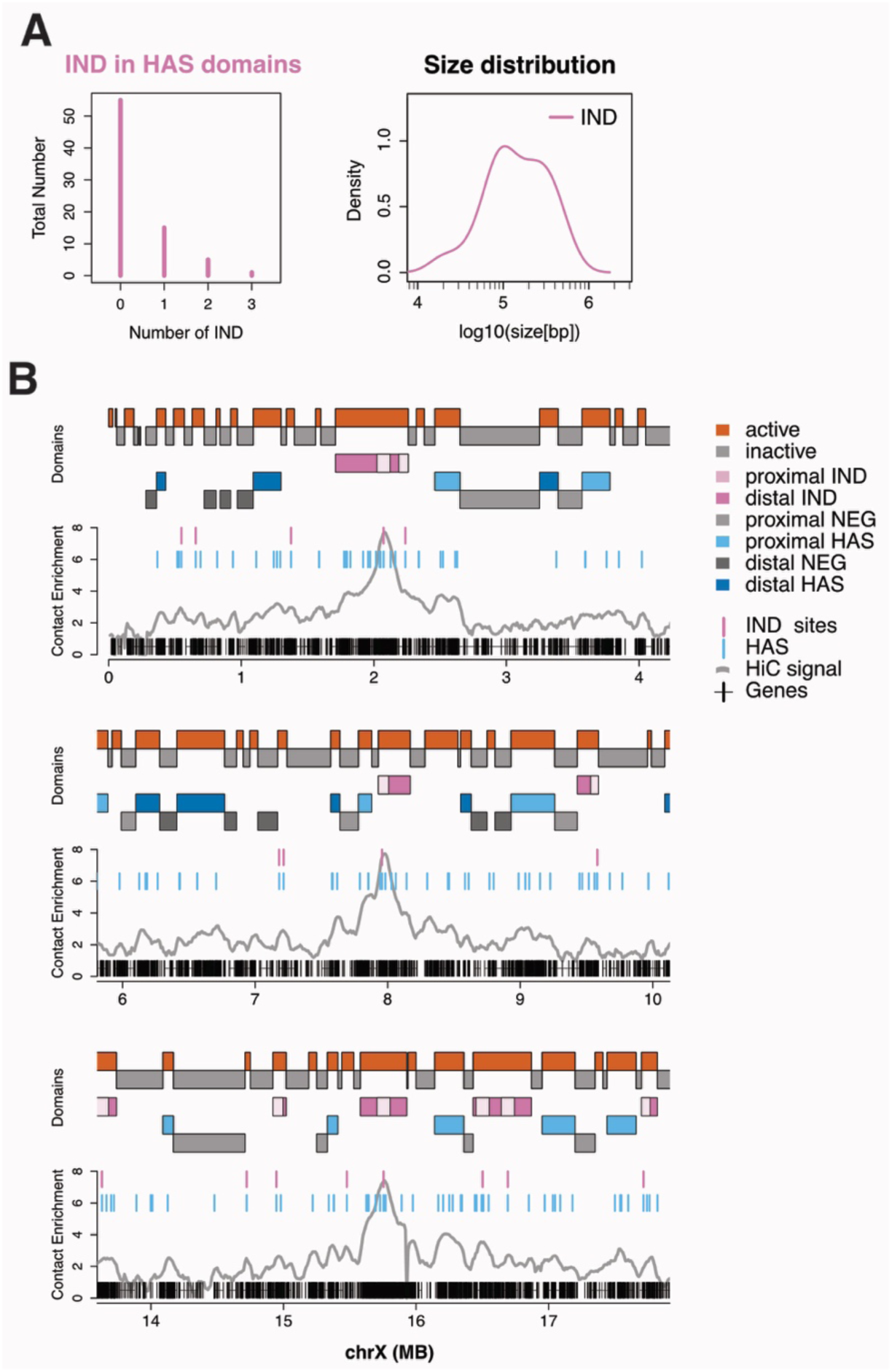
Characteristics of domains overlapping SXL RNAi induced MSL2 binding sites. **A**) *Left*: Number of induced MSL2 binding sites (IND) within domains containing HAS in the active compartment. *Right*: Size distribution of domains overlapping induced MSL2 binding sites. **B**) Similar plot as Figure 5B showing additional example regions.

## Supplementary File Legends

Supplementary File 1. Read statistics of the Hi-C data. The table indicates the number of raw reads and the number of uniquely aligned reads in each sequencing lane, the number of tags in the tag directory, the number of filtered tags in the tag directory (generated by Homer (Heinz et al. 2010)) for each sample as well as for the pooled samples.

Supplementary File 2. PC1 values for chrX and chr2L in each sample as well as for pooled samples.

Supplementary File 3. Primer sequences used for the 4C-seq library preparation.

Supplementary File 4. Read statistics of the 4C-seq data. The table indicates the number of raw reads, mapped reads, mapped X chromosomal reads and the number of reads that were filtered (non-digested, re-ligated, self-ligated) for each viewpoint, condition and replicate.

Supplementary File 5. Summary tables of the 4C-seq interaction (peak) calling in S2 control (GFP RNAi) treated cells. The first table contains peaks for each viewpoint called at various settings included in the analysis (smoothing window, minimum number of counts and z-score threshold). The second table contains the pool of interactions for each viewpoint.

Supplementary File 6. Differential analysis of the 4C-seq profiles. The first table contains summary statistics (obtained by DESeq) for all 4C fragments whereas the second for the interaction (peak) fragments.

Supplementary File 7. Genomic coordinates of SXL RNAi induced MSL-2 ChIP-seq sites in Kc cells (Villa et al. 2016).

